# Contribution of private game ranching and captive bred operations in South Africa to White rhino *Ceratotherium simum* species survival conservation

**DOI:** 10.1101/2022.09.21.508862

**Authors:** Deon Furstenburg, Michelle Otto, Pieter Van Niekerk, Derek Lewitton

## Abstract

The southern white rhinoceros (SWR) *Ceratotherium simum simum,* like all extant rhinoceros’ species, is under global threat of extinction, due to their dwindling wild population numbers in protected areas and state-owned parks as a result of poaching. The Kruger National Park (KNP), the renowned state-owned stronghold for SWR, has suffered an estimated decline of over 75.0% of its population since 2011, with the highest annual poaching rates over the past decade and a remaining population estimated at only 2,607 animals by the end of 2020, which is an average annual population decline of −10.2% from 2008 to 2020, and 2,458 animals left by the end of June 2022. On the contrary, SWR under private custodianship and management on rewilded agro-sustainable biodiversity game ranches in South Africa (estimated currently at >8,000 animals, some of which are registered Captive Breeding Operations [CBO]; assessed CBOs contained 2,882 rhinos at the time of this study (Sep 2021), have increased in population numbers and survival rates, sustaining average annual population growth performances of 9.0%. This increase has been attributed to effective security, provision of additional habitat, dispersal, and frequent genetic exchange of rhinos between breeding subpopulations by the private sector. The conservation success of the private sector has largely been overlooked and disregarded by world conservation bodies and organizations, mostly due to misguided and prejudiced media publicity and the lack of scientific analytical assessment. Private rhino custodians and their biological/ conservation breeding practices, with private agro-sustainable biodiversity wildlife management and/ or captive breeding, generally being perceived and branded as either “canned” or equated to “captive zoological-gardens”. Since the commencement of the International Convention on Biodiversity, global controversy exists whereby most of mankind today perceive bio-conservation of a species to be assigned in principle solely to protected areas and state-owned parks. The unique and advantageous roles of rewilded bio-conservation and sustainable-use conservation CBOs, being a key to green-economy and natural capital in a post-Covid-19 struggle, are mostly ignored. This study serves to assess and quantify the impact of private wildlife ranching in South Africa with specific focus on its potential contribution to rhino conservation specifically for that of the SWR *C.s. simum*.

**Data Availability Statement:** The data belong to third parties, six different private owners respectively, and cannot be shared publicly. Interested, qualified researchers can request the data by contacting the author, subject to owner’s permission. The authors confirm that they did not have any special access privileges to the data and data will be made available to other researchers in the same manner that the authors accessed it.

## Introduction

### Rhino population history

The southern white rhinoceros (SWR) is listed as Near Threatened on the IUCN red list of threatened species, but the species’ integrity and current survival status is in question, with the remaining South African SWR population in state-protected areas continuing to decrease annually. Their questionable status is propagated by noted discrepancies in available literature, reports by the IUCN and the media as well as the failure by management authorities to release up-to-date and accurate figures [1–3]. Globally, white rhino numbers have declined from >250,000 *c*.1700s, to the brink of global extinction by *c*. 1895, a single population remaining of between 20–50 individuals in what is known today as Hluhluwe-iMfolozi Park (HiMP) in South Africa [4–10]. It was protectionist and conservation practices and Project Rhino, spearheaded by the late Dr Ian Player [11], that saved the SWR from extinction, despite a genetic bottleneck that threatened their survival, status, and species integrity. Numbers in Africa increased to an estimated maximum of around 20,000 by 2011 [4,6]. Since 2011 a continuous decline, and a reduction of 75.0% reported for the largest remaining SWR population of the Kruger National Park (KNP) [1].

### Conservation shift

In South Africa, home to more than 86.5% of the world’s remaining SWRs [3,12–13], a sequence of consequential shifts of approach in SWR conservation occurred since the *c*. 1895 genetic bottleneck [6]. First priority was the protection and stabilization of the last few remaining rhinos into the establishment of a state-owned park, the HiMP of today. The second approach, to redistribute rhino across a greater geographic range and variety of habitats commenced in 1961 with the first rhino relocation to Loskop Dam Nature Reserve, and later in October the same year to the KNP. A narrative shift came in July and August 1976 when a white rhino bull and cow were introduced to the Ubizane Game Ranch of Mr. Norman Dean in KwaZulu-Natal, the first practical private ownership of SWR. Private ownership escalated since 1979 when surplus white rhinos from state-owned protected areas were sold for the first time on live game auctions. The next conservation shift happened in 1991 when the Game Theft Act of South Africa was legislated, which makes provision for legal private ownership of wildlife. This unusual situation, where a wildlife species is no longer a *res nullius* property but can be fully privately owned, has had significant consequences for SWR [14]. Legal ownership created the incentive for viable commercial trade, and consequently private ranching and breeding, and potential increase in SWR population numbers. Trade attributed also to further expansion of the specie’s distribution range. Metapopulation rhino management from private owners served as seed populations to re-establish populations throughout the SADC region as well as approved destinations globally. Consequently, the conservation status of SWR improved and in 1994, the South African SWR population was down listed to Appendix II by the Convention on International Trade in Endangered Species of Wild Fauna and Flora (CITES) for the exclusive purposes of allowing trade in live exports and hunting trophies to approved and acceptable destinations.

The present conservation approach (assessed by this study), came in 2006 with the Captive Breeding Operation (CBO) guidelines presented by the South African Development Community (SADC) Rhino Program [15], are aimed at maximizing population growth rates, ensuring long term genetic and demographic viability, animal welfare and safety, and rewilded agro-sustainable biodiversity enhancement, in specific of SWR. The first privately owned and registered SWR CBO commenced in 2008.

The IUCN World Congress (2012) adopted Motion 26 (as cited by the Department of Forestry, Fisheries and Environment [DFFE] in their Rhino Issue Management Report [16], which encourages rapid growth with genetic and demographic viability as the cornerstone goals for sustainable conservation of the species. Additionally captive populations can act as a “safety net” should the depredations of poachers reduce global “wild” rhino numbers to dangerously low levels [17]. Successful breeding programs in captivity require scientific and co-operative management to produce viable populations. Adequate husbandry, veterinary care, genetic management and veld and habitat resources management can achieve viability of captive populations [18–19]. Rhino population performance is density dependent [4] thus, distribution of localized wild populations to CBOs and private land across additional viable habitat throughout the country can maintain productive densities in the donor populations and provide founder animals to new subpopulations to enhance specie’s growth [20]. In addition, by distributing rhino geographically the possibility of enhancement of genetic heterozygosity increases [17].

Southern white rhino populations now occur widespread across South Africa in formally proclaimed conservation areas as well as on private land. The Private sector contributed approximately 2.2 million hectares of land towards rhino conservation (COP16i-38, [21]) in 2013 and increasing. As of 2020, private owners are custodians to a total of >7,000 or >57.0% of the national SWR population [22–**Error! Reference source not found.**].

### Southern white rhino (SWR) population status

According to a survey by the SADC countries Rhino Management Group and data from IUCN’s Species Survival Commission (SSC) African Rhino Specialist Group (AfRSG) [13], the last official published figures for the Southern white rhino population in South Africa was estimated at *n* = 15,625 (90.0% bootstrapped confidence index [CI]) of an estimated African SWR population of *n* = 18,067 (*n* = 17,212–18,915: 90.0% CI). A review of the historic published southern white rhino population numbers by the IUCN (including revisions) and the relevant statistical poaching trends over the past decade to date, indicate that the likelihood that the South African SWR population of *n* = 15,625 reported for December 2017 remained constant, is improbable, and most likely an overestimation by the end of 2020. From January 2008 to June 2022, a total of *n* = 9,597 rhino was recorded to have been poached in South Africa alone.

For the purpose of this report, a working estimation of the South African SWR population as of the end of 2020 was rationalized using the estimated population of *n* = 18,993 in 2012 as a starting point [24]. Given the above estimated annual white rhino losses due to poaching, and using the cautiously optimistic assumption that the national metapopulation showed a positive annual net population growth rate of between 1.0–5.0% since 2012 after poaching losses (recommended minimum goal as per the South African White Rhino Biodiversity Management Plan (BMP) for 2015–2020 [25]), this growth rate reflects the underlying biological growth rate of the metapopulation i.e. is independent of any increases or decreases in numbers due to import or export of rhino in and out of the country, this would equate to an estimated maximum national South African SWR population number of (*x*) = 14,009 by the end of 2019. Even when using an optimistic annual growth rate of 5.0%, this working estimation falls well below the short-term 5-year conservation target set by the Biodiversity Management Plan of an expected *n* = 20,400 white rhino in South Africa by the end of 2020 (which only used an estimated 2.0% annual growth rate after poaching [25]). It is also worth mentioning that South Africa suffered a severe drought in 2015–2017 along with the lag effect, decreasing the underlying growth potential as well [26].

Putatively, it has been acknowledged that the level of poaching has exceeded the per annum net births (natural underlying growth rate) and that the South African national metapopulation has been in negative growth for the past couple of years [25], and that despite poaching levels decreasing in the Kruger National Park, the numbers continue to decrease indicating that in relative terms, the percentage of this population being poached is still unsustainable (in other words, the level of poaching exceeds the per annum net births and outcompetes the underlying natural growth rate) indicating that due to continued unsustainable poaching losses, the global SWR number continues to decline.

Considering the uncertainties surrounding the true current status of the SWR, it is important to consider a range of possible conservation strategy options available in order to improve management decision making. This report serves to compare the vital rates, survival efficiency and fertility rates of a number of different options available, e.g., private ownership CBOs and free roaming state-owned protected areas.

## Materials and methods

The aim of the study is to analyse the survival and vital rate performances of a series of six (referred to as Case studies no’s 1–6) privately ranched white rhino CBOs in South Africa, to give an analytical review of the private sector and whether their management of their rhino populations has a potential detrimental effect on South African SWR survival. Secondly, to compare with the largest state-protected “wild” SWR population the KNP.

The CBOs of the Case studies are located in the Northwest, Northern Cape, Limpopo, and Free State Provinces of South Africa. Relevant data and findings of the resident rhino populations at each Case study as from date of inception to end of September 2021, as well as additional data provided by the Private Rhino Owners Association (PROA). All data provided by the private rhino owners was given willingly, without financial incentives and to the best of the corresponding authors’ knowledge, is accurate.

Natural resources, management, animal studbooks, population dynamics, and animal health and performances of the CBOs were assessed. The outcome further discussed will be in relation to the claims from the High-Level Panel (HLP) draft report of DFFE [27]) towards the conservation and survival of the species. For security purposes, the geographics, owner and property identities of the study sites are not disclosed in this report.

Using the working model depicted in Table 1 below, the working SWR population in South Africa as of the end of 2020 used for the purposes of this report, is therefore *n* = 14,035~14,000, but most likely less at the time of publication due to the additional *n* = 451 poached in 2021 and *n* = 259 rhino lost already up to the end of June 2022 (or *n* = 16,225 of remaining African rhino extrapolated number at 86.5%, or *n* = 16,339 remaining global SWR number extrapolated at 99.3%). This working South African white rhino population number calculated in this study, is very similar to an independent recent publication of an estimated South African SWR number of *n* = 14,410 for 2020 [1].

**Table 1.**
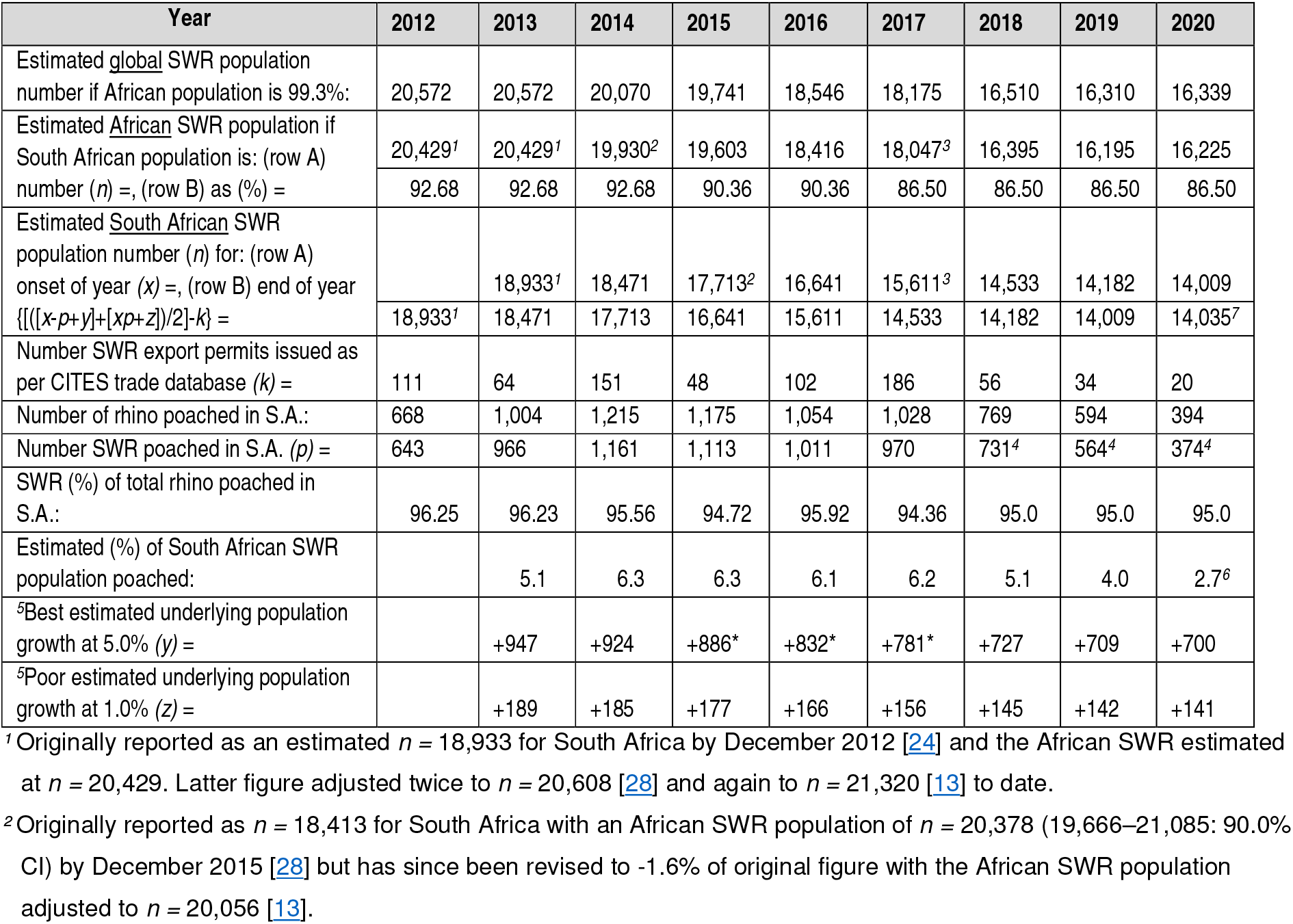
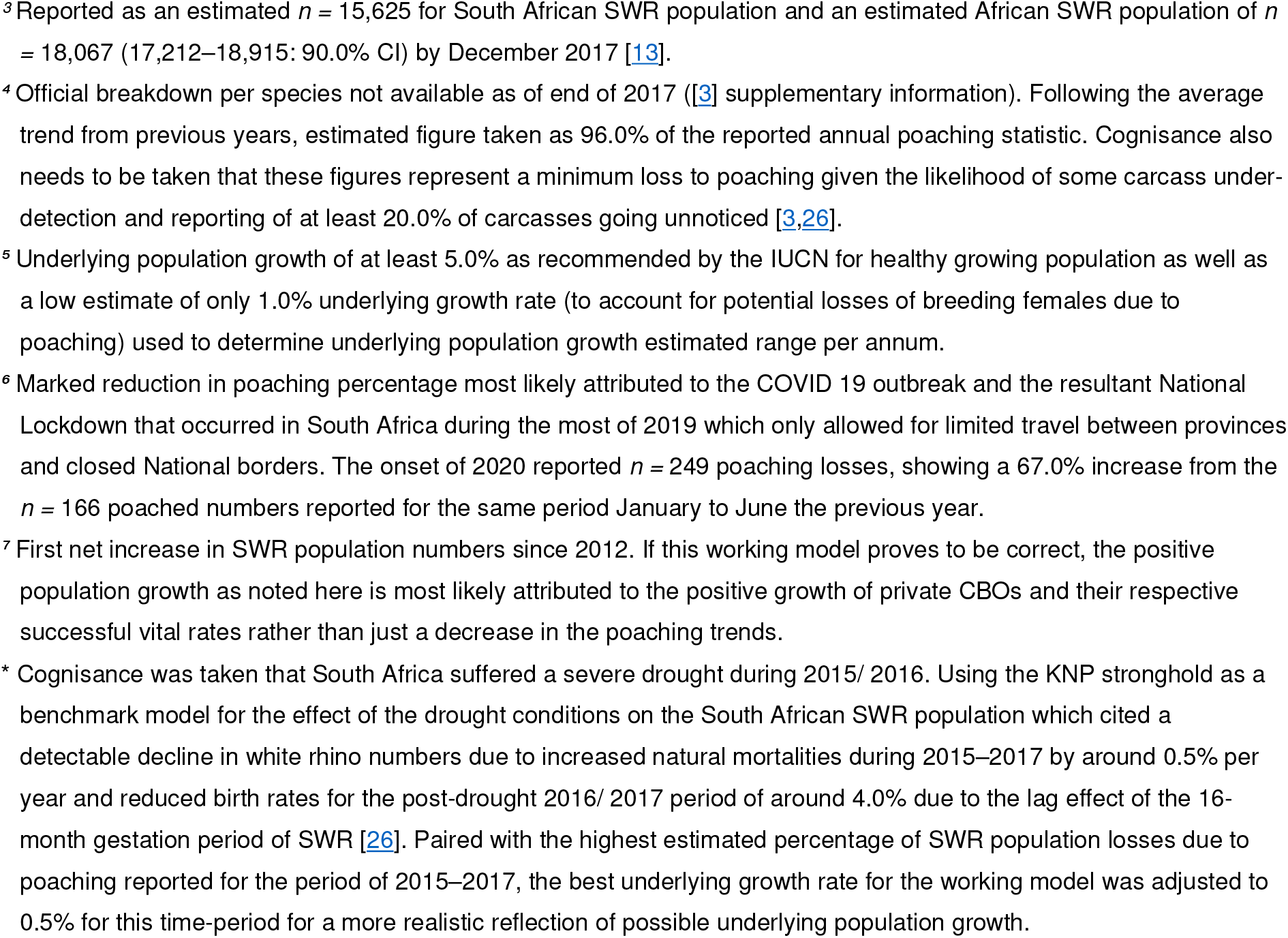
Model of working South African southern white rhino (SWR) population estimates.

### Operation case studies (study area)

#### Case study no. 1

The CBO commenced in March 2008, was officially registered as a Captive Breeding Operation for Southern white rhino in May 2009 and had a resident SWR population distributed into *n* = 21 subpopulations of *n* = 2,000 rhinos at end of May 2021 (Table 2), which contributes to >14.0% of South Africa’s estimated working national population number by the end of 2020. The period reviewed is approximately 14 years. Environment is the Dry Highveld Grassland Bioregion in Vaal-Vet Sandy Grassland with pockets of Klerksdorp Thornveld [29], consisting of 8,516 ha of land inhabited by rhino. Some 36.8% of this land had been rewilded from historically ploughed crop lands and 59.9% from past livestock farming: 96.7% of the land now sharing a biodiverse multispecies mixture of natural game and fowl being conserved.

**Table 2.** Geographic, spatial and management parameters of the white rhinoceros captive bred operations studied.

| Parameter | CBO Case study no. |  |  |  |  |  | Total / Average |
| --- | --- | --- | --- | --- | --- | --- | --- |
|  | 1 | 2 | 3 | 4 | 5 | 6 |  |
| Registration: | May 2009 | Sep 2019 | Feb 2021 | Feb 2021 | In process | 2022 |  |
| Operation starting date: | Mar 2008 | Aug 2013 | 2013 | 2011 | 2011 | 2009 | 11 Years |
| Geographic location (Province): | North West | Northern Cape | Limpopo | Limpopo | Limpopo | Free State | 4 Provinces |
| Total rhino land area size (ha) = | 8,516 | 7,955 | 2,502 | 2,686 | 1,110 | 2,939 | 25,708 |
| Area of land rewilded from ploughed land: (row A) per (ha) =, | 3,135 | 0 | 5 | 105 | 500 | 1,064 | 4,809 |
| (row B) as (%) = | 36.8 | 0.0 | 0.2 | 3.9 | 45.1 | 36.2 | 20.4 |
| Area of land rewilded from alien livestock farming: (row A) per (ha) =, | 5,101 | 7,788 | 2,500 | 2,571 | 514 | 2,874 | 21,348 |
| (row B) as (%) = | 59.9 | 97.9 | 99.9 | 95.7 | 46.3 | 97.8 | 82.9 |
| Total rhino land area rewilded (%): | 96.7 | 97.9 | 100.0 | 99.6 | 91.4 | 100.0 | 97.6 |
| Rhino total population size (*n*) = | 2,000 | 306 | 145 | 200 | 78 | 153 | 2,882 |
| Rhino population as (%) of current South Africa = | 14.3 | 2.2 | 1.0 | 1.4 | 0.6 | 1.1 | 20.6 |
| Number of rhino herds/ subpopulations in system (*n*) = | 21 | 6 | 2 | 4 | 1 | 7 | 41 |
| Average number of rhino (*n*) per herd [population density] = | 95 | 51 | 73 | 50 | 78 | 22 | 61 |
| Range size per subpopulation (min-max range, ha) = | 100–594 | 750–3,200 | 890–1,605 | 569–806 | 877 | 282–496 | 100–1,605 |
| Average rhino range size (ha) = | 393 | 944 | 1,251 | 663 | 877 | 367 | 749 |
| Median (middle value) range size (ha) = | 426 | 1,029 | 1,251 | 614 | 877 | 351 | 758 |
| Animal stocking density (min-max range, ha/rhino) = | 3.2–13.7 | 13.8–133.3 | 13.4–35.9 | 8.3–25.2 | 11.2 | 14.8–51.0 | 3.2–133.3 |
| Average stocking density (ha/rhino) = | 4.3 | 26.0 | 17.3 | 13.4 | 14.2 | 19.2 | 15.7 |
| Veld management strategy per rhino subpopulation: | 2-sectional rotational animal stocking | 4-sectional rotational animal stocking | 2 single systems, multiple fire/ burn blocks | 4 single systems, multiple fire/ burn blocks | 1 single system, multiple fire/ burn blocks | 7 single systems, multiple fire/ burn blocks |  |
| Mean annual long-term rainfall (mm) = | 689 | 300 | 507 | 602 | 454 | 573 | 300–689 |

#### Case study no. 2

The CBO commenced in August 2013, was officially registered as a CBO in August 2019 and includes *n* = 5 white rhino subpopulations which totalled *n* = 306 rhinos at the end of May 2021 (Table 2). Review period is approximately 8 years. Environment is the Eastern Kalahari Bushveld Bioregion in Olifantshoek Plains Thornveld, with pockets of Kuruman Mountain Bushveld and Northern Upper Karoo [29], consisting of 7,955 ha of land inhabited by rhino. A total of 97.9% of this land had been rewilded from monoculture livestock farming to a biodiverse multispecies (*n* = 10) mixture of natural plains game and conserved.

#### Case study no. 3

The CBO commenced in 2013, was officially registered as a CBO in February 2021 and includes *n* = 2 white rhino subpopulations which totalled *n* = 145 rhinos at the end of July 2021 (Table 2). Review period is approximately 8 years. Environment is the Sub-Escarpment Savannah Bioregion in Granite Lowveld, with elements of Phalaborwa-Timbavati Mopaneveld [29], consisting of 2,502 ha of land inhabited by rhino of which 99.9% had been rewilded from monoculture livestock farming to a biodiverse multispecies (*n* = 14) mixture of natural plains game and conserved.

#### Case study no. 4

The CBO commenced in 2011, was officially registered as a CBO in February 2021 and includes *n* = 4 white rhino subpopulations which totalled *n* = 200 rhinos at the end of July 2021 (Table 2). Review period is approximately 10 years. Environment is the Sub-Escarpment Savannah Bioregion in Granite Lowveld, with elements of Phalaborwa-Timbavati Mopaneveld [29], consisting of 2,686 ha of land inhabited by rhino of which 3.9% had been rewilded from historically ploughed agricultural lands and 95.7% had been rewilded from historically monoculture livestock farming to a biodiverse multispecies (*n* = 13) mixture of natural plains game and conserved.

#### Case study no. 5

The CBO commenced in 2011, official registration as a CBO in progress at time of this study and includes *n* = 4 white rhino subpopulations which totalled *n* = 78 rhinos at the end of July 2021 (Table 2). Review period is approximately 10 years. Environment is the Central Bushveld Bioregion in Mkhado Sweet Bushveld [29], consisting of 1,110 ha of land inhabited by rhino of which 45.0% had been rewilded from historically ploughed agricultural lands and 46.3% had been rewilded from historically monoculture livestock farming to a biodiverse multispecies (*n* = 14) mixture of natural plains game and conserved.

#### Case study no. 6

The CBO commenced in 2009, was registered as a CBO in March 2021 and includes *n* = 7 white rhino subpopulations which totalled *n* = 153 rhinos at the end of November 2021 (Table 2). Review period is approximately 12 years. Environment is the Dry Highveld Grassland Bioregion of SA in both Vaal-Vet Sandy Grassland and the Central Free State Grassland [29], consisting of 2,939 ha inhabited by rhino of which 37.8% had been rewilded from historically ploughed agricultural lands and 46.3% had been rewilded from historically monoculture livestock farming to a biodiverse multispecies (*n* = 12) mixture of natural plains game and conserved.

### Statistics

Average annual natural mortality rates (M_t_) was calculated using the overall known naturally mortality (M) figures divided by the net population {[Introductions (I), plus births (B) minus removals (R)] over the period (t) of production at each Case study site}:

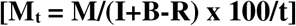

Similarly, the average annual poaching rate (P_t_) was calculated using the known overall poaching (P) figures divided by the net population [Introductions (I), plus births (B) minus removals (R) minus natural mortality losses (M)] over the period (t) of production at each Case study site:

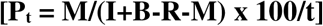

The average annual calf recruitment birth rate (B_t_) was calculated using the known per annum births divided by the number of adult females (F) present in each population at the onset of the year over the period (t) of production at each Case study site:

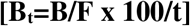

The average annual growth rate (Gt) was calculated using the annual biological growth rate [annual births (B) minus the annual natural mortalities (M)] achieved per annum divided by the population figure present at the onset of the year (Po) over the period (t) of production at each Case study site:

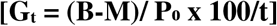

## Results

### Geographic scope of rhino operations reviewed

The distribution of the white rhino subpopulations in South Africa is fragmented, all existing in fenced-in, protected areas either in state-owned national parks managed by South African National (SAN)Parks and reserves managed by provincial conservation authorities, or non-state areas comprising of private/ community game reserves, and/ or private ranches, with white rhino regarded as a common species [19,30]. According to DFFE [31] the total area occupied by SWR within South Africa in 2015 exceeded 49,000 km^2^ (4.9 Mha), of which approximately 18,000 km^2^ (1.8 Mha) was private or communal land. The considered “wild” (though fenced-in) SWR numbers were estimated at *n* = 17,208 by 2015, of which *n* = 12,273 (72.0%) occurred on state-owned land, and *n* = 4,735 (28.0%) on private land, whilst the KNP subpopulation was estimated at *n* = 8,365–9,337 (Fig 1). In 2020 the HLP draft report and the IUCN recognised a global population size of *n* = 10,080 mature animals remaining [3] which included privately ranched populations reported at an estimated modelled (R^2^ =0.988) total size of *n* >8,000 in 2022 (Fig 2), thus raising concerns regarding the proposed motions by DFFE with regards to the potential detrimental effect on green or natural capital and biodiversity conservation by the private sector since the private sector now owns more white rhinos than the entire of the protected areas and the rest of Africa accumulative. The positive slight upward turn in numbers of the South African and global SWR populations in 2020 (Table 1 and Fig 1) are being attributed to the growth success of SWR populations on private land (CBOs).

**Fig 1.**
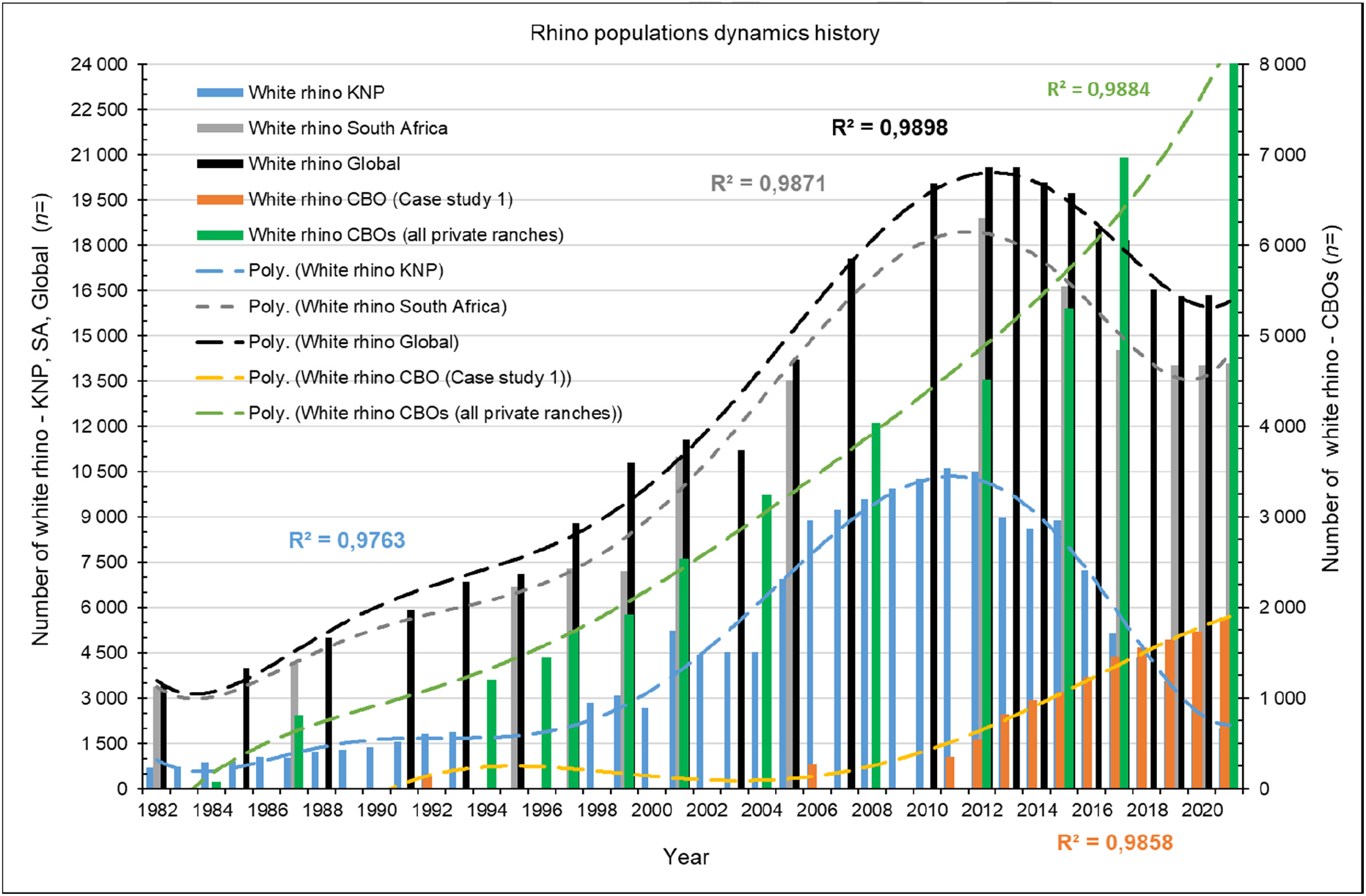
Metapopulation dynamics of southern white rhino numbers since 1982, respectively for SA, KNP, all private ranches accumulative, and for CBO Case study no. 1. Data included from [1,3,6,12,22–**Error! Reference source not found.**,25,32–37].

**Fig 2.**
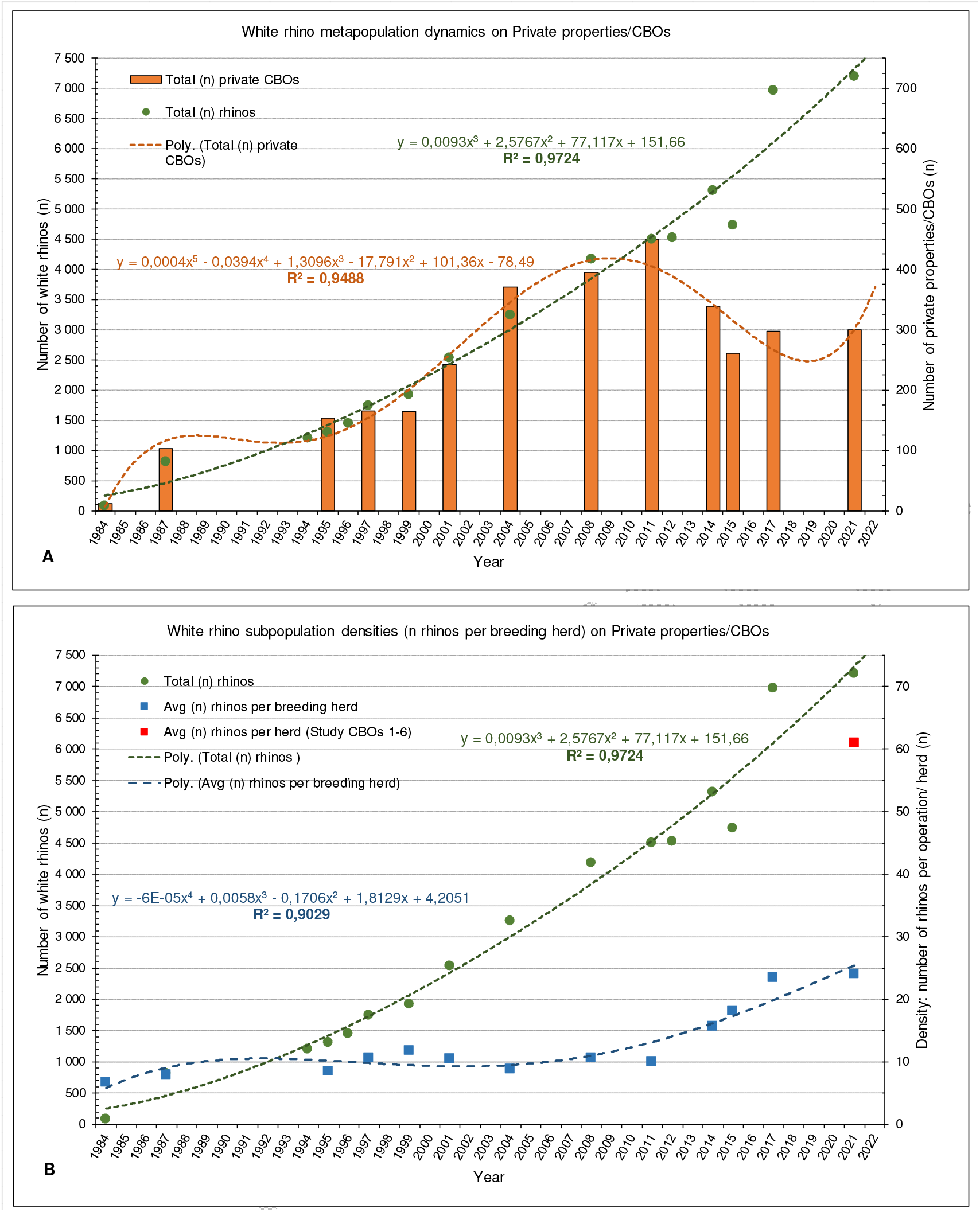
Annual southern white rhino metapopulation dynamics [rhino numbers (*n*)] on private properties and CBOs across SA. (A) as per number of ranches/ subpopulations (*n*), (B) average animal density/ number of rhino (*n*) per CBO subpopulation/ breeding herd, with 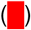 accumulative average density/ number of rhino (*n*) for Case studies no’s 1–6. Data included also from [12,**Error! Reference source not found.**,**Error! Reference source not found.**,38–40].

A trend of steady increase exists in the size of the privately owned populations since 1980 (Fig 2). Response rate of questionares successfully retrieved from private rhino owners with the 2015 survey [12] was 28.7% (that is 75/261, representing 1,669,000 ha of land inhabited by *n* = 5,225 rhino) as per the PROA database of the number of private/ communal properties being identified to have white rhino. Response rate for the 2017 survey was 25.3% (75/297), representing 959,000 ha with *n* = 4,029 rhinos. A follow-up independent attempt by PROA to acquire the outstanding rhino numbers resulted in an estimate of *n* = 6,968 [12], a difference of *n* = 2,940 rhinos between the two estimates for 2017. Private owners are withholding information due to the exponential rise in poaching since the early 2000s (Fig 3). It can be accepted with confidence that the number of rhinos in private ownership recorded with the surveys as from 2004 and after are underestimated, the population be greater than shown in Fig 1 and Fig 2. If one assumes an average growth of 6.0% per annum off the 2014 base of *n* = 5,225 white rhino, that will result in *n* = 6,218 animals on private and communal property, it suggests that *n* = 2,189 (or 35.0%) of the rhinos were not reported in the 2017 survey.

**Fig 3.**
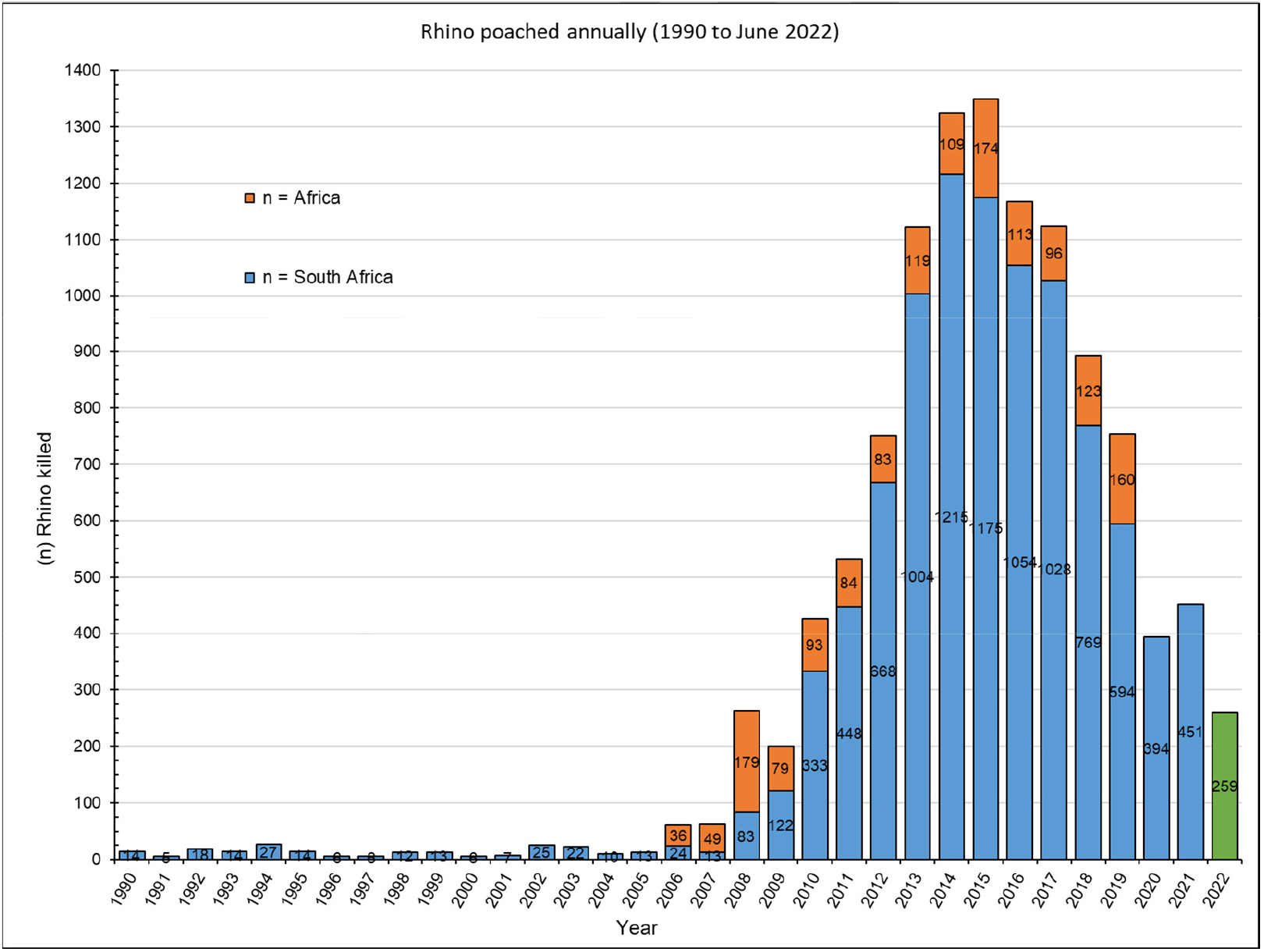
History of recorded number of rhinos poached per annum (2022 for 6 months to end June). Data compiled from [1,5,13,41–42].

A significant decline in the number of privately managed white rhinos in Limpopo Province from 2008 to 2020 (Fig 4) can be ascribed to the escalating risk to poaching, 46.9% (174/371) of all recorded incidents across country occurred in Limpopo. Also, the national decline in number of private rhino properties can be ascribed in three-fold to escalating poaching risk, increased security management costs, and the ban on horn-trade to generate income for covering operational costs. The increase of rhino numbers in Northwest and Northern Cape is a dual consequence of less poaching risk (10.5% and 0.8% respectively) and the high-performance annual growth rates of Case study no’s 1 & 2 in accumulation inhabiting more than *n* = 2,250 rhinos at the time.

**Fig 4.**
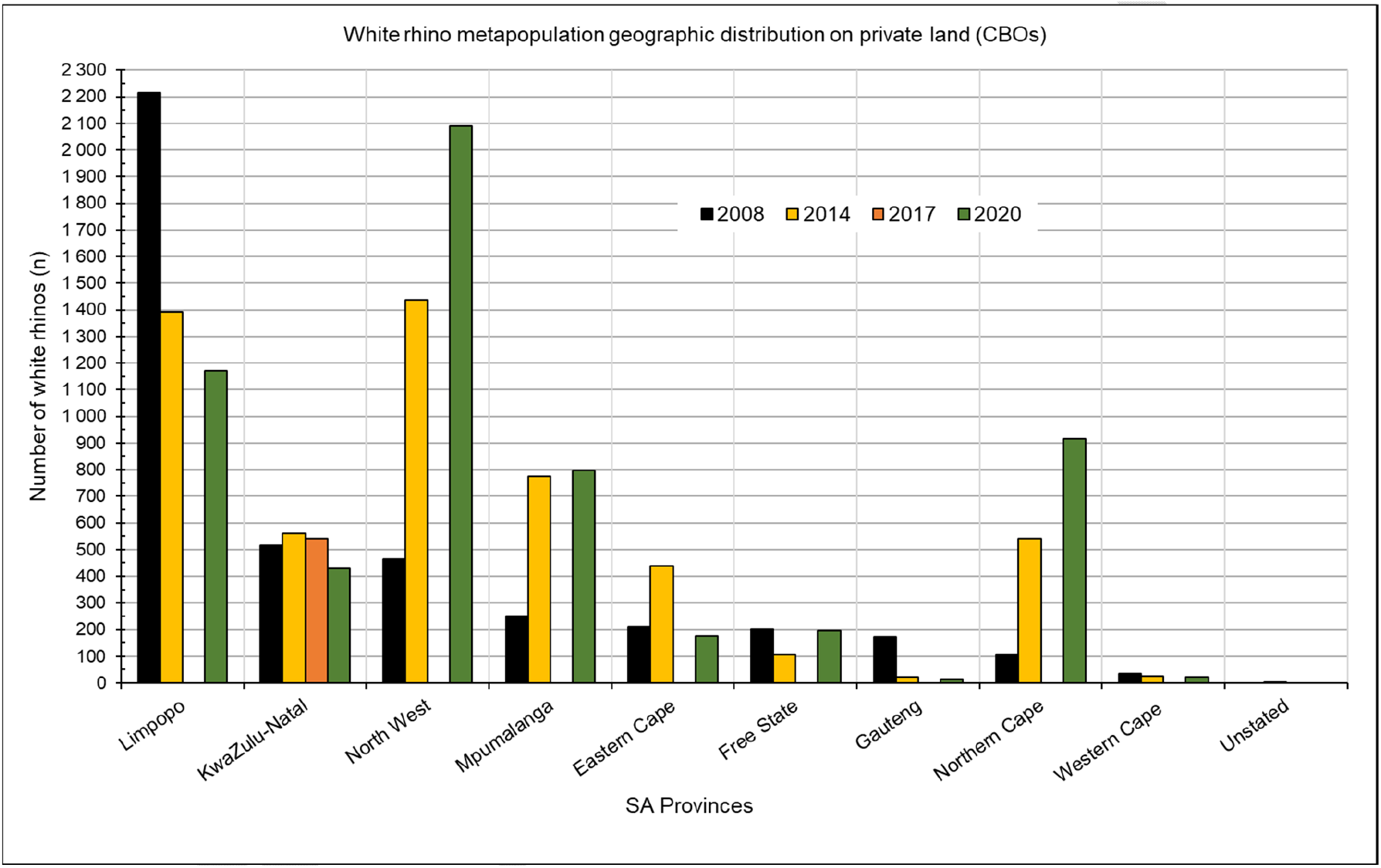
Metapopulation geographic distribution of SWR on private land (CBOs) across SA (years 2008, 2014, 2017 and 2020). Data surveys from [12,**Error! Reference source not found.**,38,43].

Despite CBO registrations and macro-genetic metapopulation genetic diversification through gene migration/ gene flow by way of frequent translocations (exchange) and relocations [19,22,44–48] across more than 300 privately managed populations, the HLP draft report [27] claims these rhinos to be genetically inferior and of little to no value to conservation and survival of the species. Such claiming is in direct juxtaposition of review and reports by the IUCN African Rhino Specialists group, which states that the CBOs of Case study no.1 is recognized as a **Key 1** population, vital to the continued conservation of South Africa’s SWR population.

### Operation description and intensity

#### Case study no. 1

Case study no. 1 follows the IUCN’s recommendations for conservation breeding. The operation controls certain aspects of the rhino’s environment using of a system of large (> 100 ha) camps (Fig 5), each of which is subdivided into two sister-camps to allow for the rotation of grazing in a manner that attempts to sustain the ecological condition of the veld. When grass growth exceeds consumption during moist summer months, this grass is mown and removed. During dry winter months or during drought periods the rhinos are provided with supplementary feed in demarcated locations. Supplementary feeding can make up to 48.0% of the rhino’s daily dietary intake during these periods. Additional feeding is designed to maintain the reproductive condition of the animals and to maintain their general well-being. Access to clean drinking water is ensured throughout the year for all rhinos. Each subpopulation contains multiple dominant breeding bulls to allow for self-choice of own mate selection, thus aligned with natural mating behaviour. General ecological requirements such as adequate shade and shelter, mud baths and rubbing posts are available. The CBO is described as a semi-intensive, semi-free roaming, rewilded agro-sustainable biodiversity wildlife management system, being more intensive (restricted) than free roaming “wild” populations, but less intensive than zoos or most safari parks worldwide (in other words, it is a semi-wild captive breeding operation) making it unique in existence.

**Fig 5.**
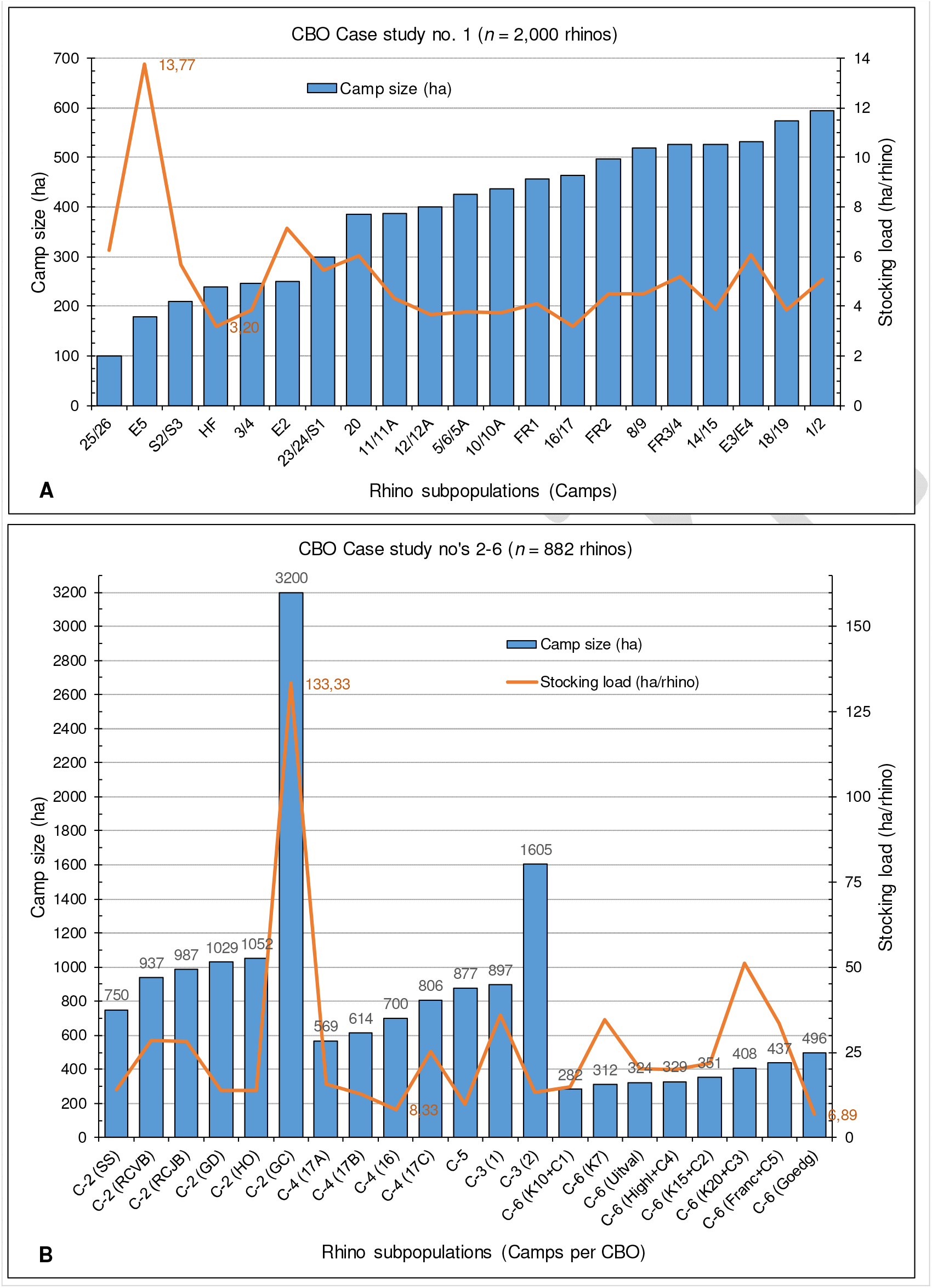
The *status quo* (2021) stocking density for each white rhino subpopulation as per private CBO studied. (A) Case study no. 1 *n* = 21 subpopulations, (B) Case studies no’s 2 (C-2), 3 (C-3), 4 (C-4), 5 (C-5), 6 (C-6), *n* = 21 subpopulations.

#### Case study no. 2

The operation is similar in its outlay, strategy and management to that of Case study no. 1, though with differences: (a) camps are generally larger in size size (Fig 5), all exceeding the minimum natural territorial range of 700 ha for the species [49]; each of which is (b) subdivided into either *n* = 2, 3 or 4 sister-camps to allow for rotational grazing – each rhino thus roams a >700 ha range; (c) all rhinos return passively to a centred 3–4 ha walk-in boma (a separate boma for each subpopulation) every night for safety – boma gates are opened in the morning when the rhinos exit on own account to the roaming fields until late afternoon when the rhinos return on own account lured by a ration of quality animal-feed pellets provided in the evening; (d) the environment is more arid with less fodder, thus daily supplementary feeding contribute up to 70.0%; and (e) the CBO also co-inhabits multiple other wild game species. The CBO is described as intensive, a limited-free roaming, rewilded agro-sustainable biodiversity wildlife management system, being more intensively managed to Case study no. 1 by way of daily feeding and nocturnal boma confinement for security reasons.

#### Case studies no’s 3 to 6

The CBOs are most similar to each other, though different to that of Case studies no’s 1 and 2. A single large semi-free roaming camp used that is not subdivided. Except for that of Case study no. 6, the subpopulation camps mostly exceed the minimum natural territorial range, and some also exceeds the minimum natural home range for the species size (Fig 5). Rotational grazing (veld-rest) accomplished by a program of rotational burning of moribund vigour, is done in a series of different block burns within each camp. Winter and drought supplement feeding are provided at random across the camp (no permanent feeding areas), thus the rhinos are more “wild” and less associated with human presence. Although the same breeding principles of Case study no. 1 are followed, lack of daily monitoring allows for less analytical data per individual rhino to be recorded. The CBOs are described as semi-extensive, semi-free roaming, rewilded or protected agro-sustainable biodiversity wildlife management systems of special uniqueness, in difference to Case studies no’s 1 and 2.

### Management parameters of CBOs

Ecological management of the habitats is also incorporated in the management plans/ BMPs of all the CBOs studied – analysis of the hydrology, climate, geology and soils, landscape, habitats and vegetation of the area. The management strategy followed allows for continual annual analysis of the natural environment (habitats, ecological veld condition, and vegetative carrying capacities) for effective continued best practice animal stocking load and environmental management.

Management decisions are always made with sustainability in mind. In some rhino camps, analysis of the vegetation determines fire regimes, if any should be followed, to keep the floristic composition productive (healthy eco-functional) and structurally viable. Managerial adjustments include the availability of different water sources (natural pans and/ or streams) throughout each camp by the addition of either water troughs and/ or the making of earth dams, depending on the geographic outlay. Each camp therefore has several natural water sources available *ad lib* to the rhino throughout the year. Built water troughs are cleaned weekly and routinely inspected.

The habitats inhabited by the resident rhino at the CBO Case studies fall within that of the native distribution of origin of the species [49–50]. The environments are controlled however as the rhino are kept in fenced-in rewilded [51] camps for their own protection and to prevent the animals from escaping. It needs to be emphasized that the major of protected areas in South Africa (including most national parks and the KNP) and referred to by government and the IUCN as “wild” populations are in fact fenced-in camps (large camps, ranging from <100 ha to 1.98 Mha for KNP), as indicated in Fig 6 and Fig 7.

**Fig 6.**
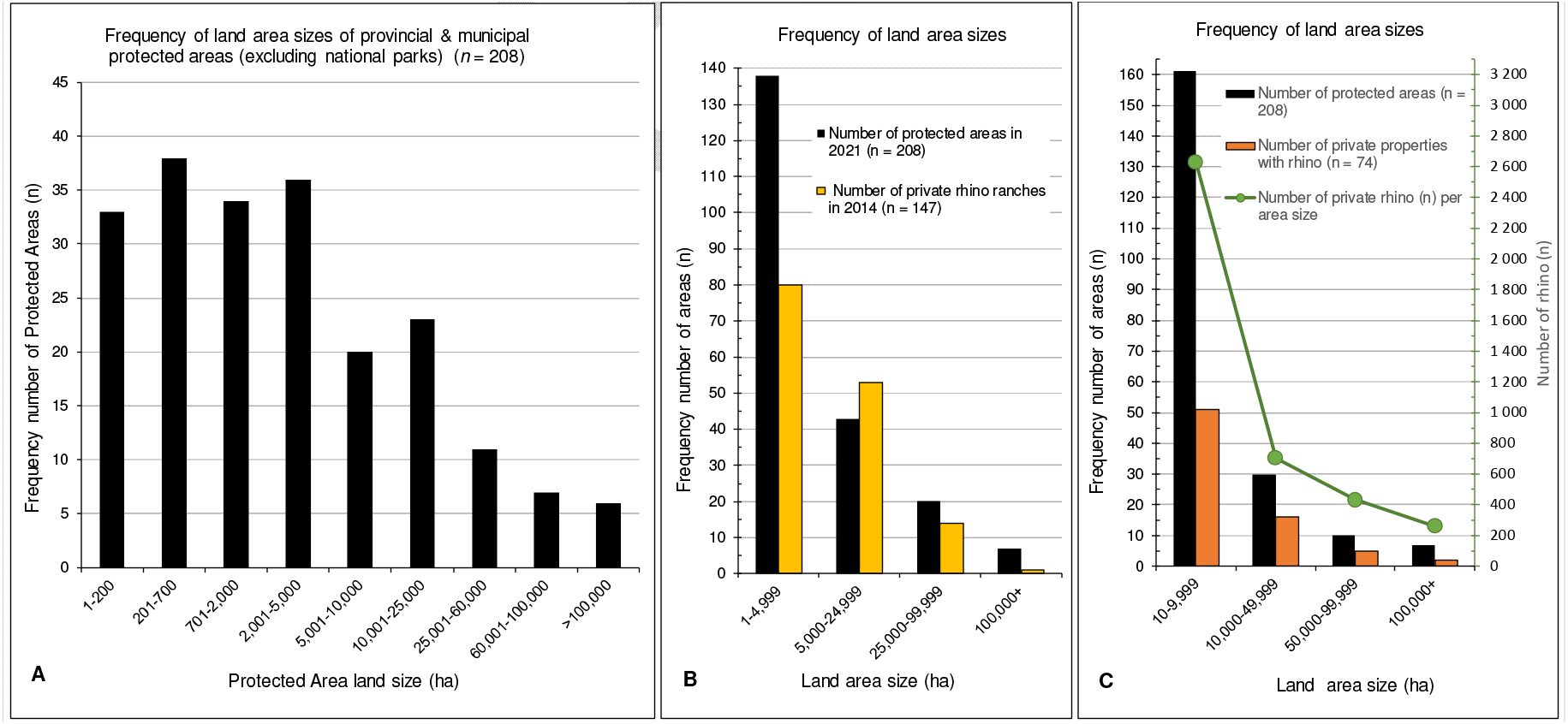
Frequency distribution ratio of land area sizes (hectares) inhabited by white rhino populations on protected areas compared to private populations. (A) Protected areas (provincial & municipal, excluding national parks), (B & C) Private properties as for 2014 and 2017, survey data from [12,**Error! Reference source not found.**,38].

**Fig 7.**
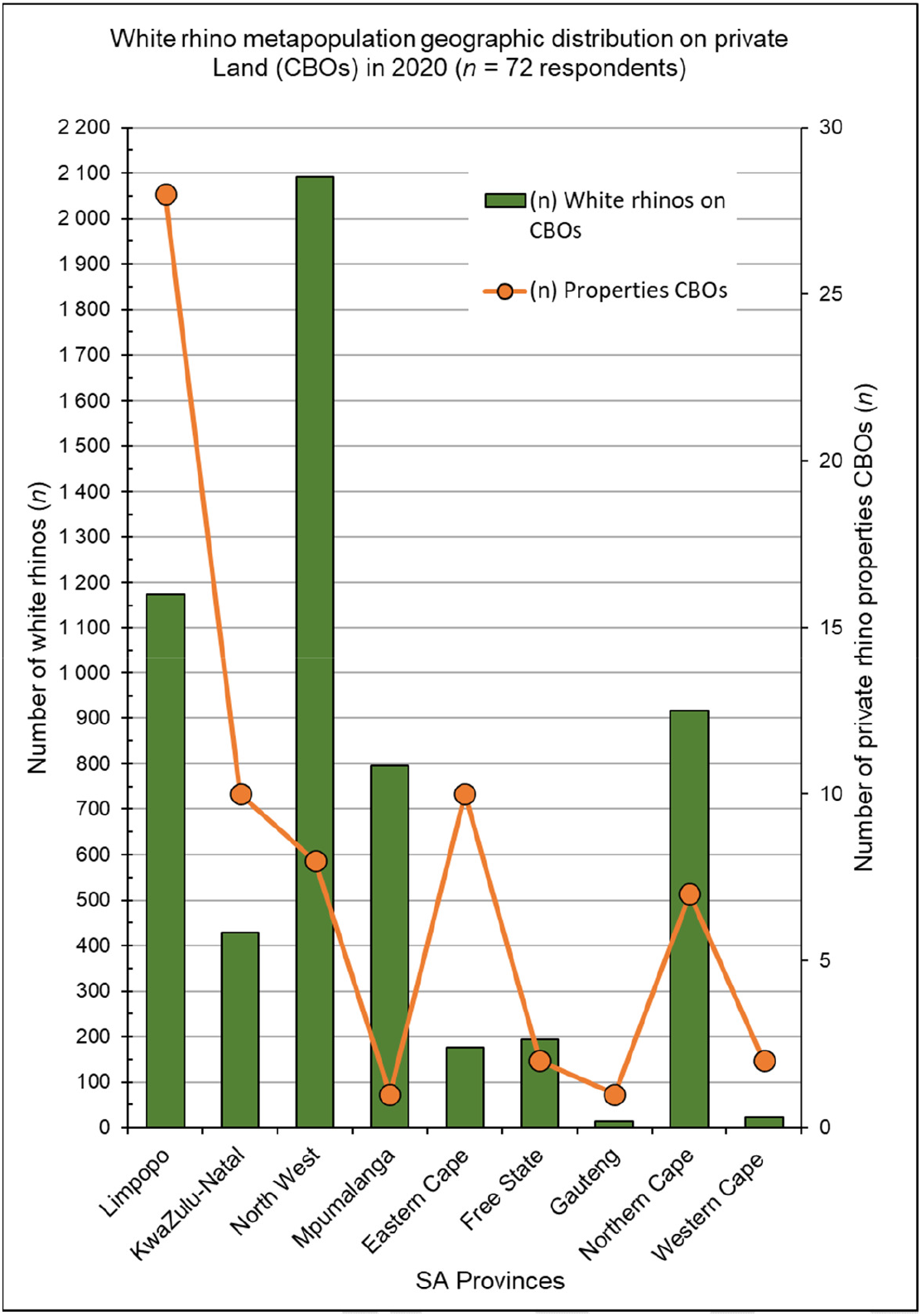
Geographic distribution ratio of an estimated 24.0% of the private white rhino metapopulation in South Africa as at end of 2020. Data adapted from [**Error! Reference source not found.**].

The breeding policy followed is to allow natural mating to occur within each subpopulation, without human intervention other than controlling the subpopulation numbers and stocking densities. To reduce numbers and potential inbreeding, selected sub-adults (2.5–4.0 years age) are removed from their natal subpopulations and either, relocated to new camps, or traded (sold) to other CBOs, game ranches or protected areas. This allows biological macro-genetic metapopulation diversification through gene migration/ gene flow [19,22,44,46–47,52] and subpopulation control, limits inbreeding and allows for further potential genetic diversification (increased genetic integrity) by establishing new mixed breeding subpopulations. Each breeding subpopulation consists out of a minimum of *n* = 2–4 dominant breeding bulls (*n* >10 for Case studies no’s 3 & 5), *n* = 20–30 or more adult females, *n* = 10–25 or more calves and juveniles and *n* = 10–25 or more sub adults. The rhinos in each breeding subpopulation are therefore allowed to create their own natural hierarchy and herd structure as in free roaming “wild” conditions.

The presence of at least two dominant bulls in each breeding subpopulation, allows breeding females to naturally select their own mate for breeding and to promote heterozygosity. In addition, an average of *n* = 4–5 or more subordinate bulls are also left in each breeding subpopulation to allow for natural hierarchy establishment and integration into the socio-dynamics of the herd. If becoming sexually active and continues to be accepted in the social structure of the herd, such a bull is allowed by management to stay and become an additional replacement breeding bull. Bull selection in each subpopulation is therefore determined by temperament, breeding ability and social integration. Strict monitoring of the performance and/ or socio-dynamics of each herd is maintained. Keeping a studbook and camp listing of each herd aid as further prevention to potential inbreeding. Due to the high reproductive rates accomplished by the CBOs, additional land is constantly bought and added for expansion, e.g., Case study no. 1 commenced with 3,455 ha in 2009 and now in extent of 8,516 ha, with added additions in 2014, 2015, 2016 and 2018, to establish new breeding herds sourced predominantly from progeny born from within the CBO.

### Genetic heterozygosity and species integrity

Genetic diversity is a very important feature of living organisms. It serves for population adapting to the environment. With increased allelic variation, individuals display adaptive characteristics that suite the environment, so genetic diversity is essential for species survival. Gene immigration/ “gene flow” is the transfer of genetic material from one population to another by migration of individuals or gametes. This can alter genetic diversity by changing allelic frequencies in a population. Gene flow is therefore very important to reduce genetic drift and effects and is extremely important for conservation genetics [53].

Today, “wild” southern white rhino occur in isolated populations within South Africa, such as the KNP, HiMP and other smaller SANParks, with limited to no gene flow between them. Furthermore, when the marked ongoing losses due to poaching at these sites are also taken into consideration, the contraction of these wild populations in the absence of gene flow from other sources, negatively affect the genetic diversity and evolutionary potential of the South African SWR through potential genetic drift. The potential for further loss of allelic diversity in these wild populations over time can however be limited by new management regimes that stimulate gene flow via periodic immigration (introduction) of new individuals, thereby preventing isolation and reducing population differentiation, as is used by the private sector.

Rhino on private land, unlike their state-owned “wild” counterparts, are frequently translocated and relocated to establish new seed populations. Translocations on private land are carried out to augment existing populations and/ or remove excess numbers from their populations as a form of population control. These excess rhinos are then sold and translocated to other, new populations at a different geographic site which promotes gene flow amongst populations, thereby reducing the potential detrimental effects of inbreeding depression [54] within the South African southern white rhino metapopulation.

The six Case studies assessed in this study showed that the geographical and genetic origin of the resident rhino populations were diverse, consisting out of rhino sourced originally from both free-roaming “wild” populations and privately managed populations. The *n* = 1,511 rhino, founder (F0) animals sourced from off the properties, have been introduced accumulatively to the six CBOs studied (Table 3), consisted out of various age classes ranging from one month old to adult, and included both sexes. Of the introductions, *n* = 1,206 (79.8%) were sourced from other private owners or game ranches and *n* = 305 (20.2%) directly from “wild” (fenced-in protected areas) populations including KNP, HiMP, other SANParks and provincial reserves. The introductions happened at periodic intervals as indicated in in Fig 8, and some more than once from the same external population. In total, rhino translocated to the studied CBOs were sourced from *n* = 176 different founder populations (genetic pools), consequently potentially contributing *n* = 176 different genetic source migrations. Within each CBO these founder members from different geographical sites were also admixed into different breeding populations, therefore further enhancing the potential gene flow in each breeding subpopulation.

**Fig 8.**
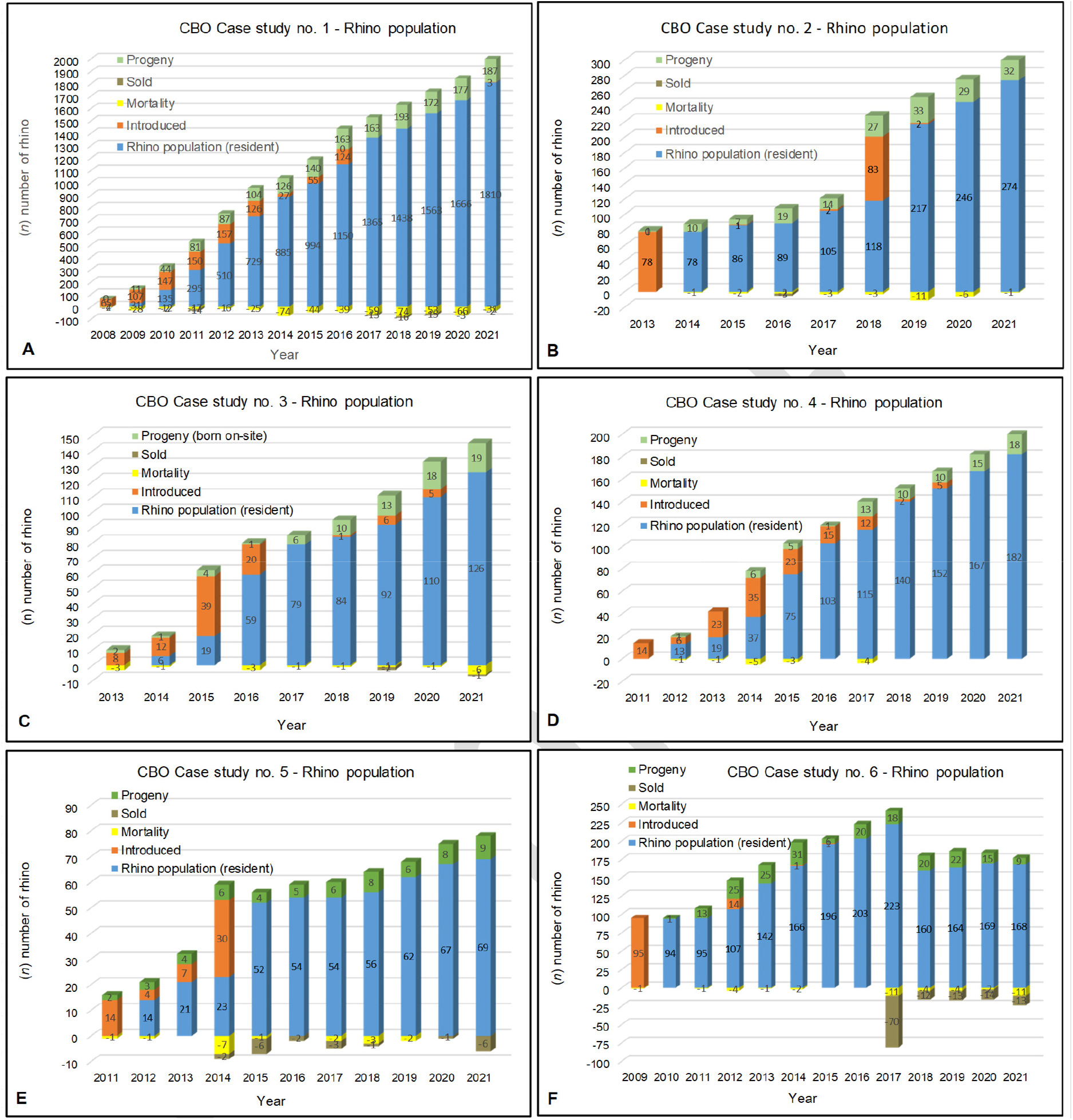
Annual population history indicating performance growth for CBO Case studies no’s 1–6.

**Table 3.** Population parameters and production vital rates of the white rhinoceros captive breeding operations studied.

| Parameter |  | CBO Case study no. |  |  |  |  |  | Total / Average | NKP |
| --- | --- | --- | --- | --- | --- | --- | --- | --- | --- |
|  |  | 1 | 2 | 3 | 4 | 5 | 6 |  |  |
| Genetic diversity potential | Number (n) of Rhino sourced from external populations: (row A) male =, (row B) female =, (row C) unknown sex = | 300 | 48 | 30 | 49 | 12 | 11 | 450 | ? |
|  |  | 661 | 118 | 65 | 74 | 43 | 12 | 973 | ? |
|  |  |  |  |  |  |  | 88 | 88 |  |
|  | Total number externally sourced (n) = | 961 | 166 | 95 | 123 | 55 | 111 | 1,511 | 351 |
|  | Percentage sourced directly from wild populations including KNP (%): | 17.7 | 68.1 | 6.0 | 3.0 | 4.0 | 52.3 | 25.2 | 100.0 |
|  | Population size at time of analysis (n) = | 2,000 | 306 | 145 | 200 | 78 | 153 | 2,882 | 2,607 |
|  | Number of different external populations founders sourced from (n) = | 95 | 19 | 11 | 24 | 9 | 20 | 178 | 1 |
|  | Genetic diversity potential: | Excellent | Good | Moderate | Good | Moderate | Good | Good | Poor |
| Vital rates | Total number of known progeny produced (n) = | 1,648 | 172 | 74 | 79 | 61 | 205 | 2,239 |  |
|  | Total number of known natural deaths (n) = | 508 | 306 | 8 | 6 | 12 | 30 | 870 |  |
|  | Average natural mortality rate (%): | 1.43 | 0.86 | 0.55 | 0.25 | 1.15 | 1.19 | 0.9 | >1.1 |
|  | Total known number poached (n) = | 32 | 3 | 9 | 8 | 5 | 11 | 68 | >5,142 |
|  | Average annual Poaching rate (%): | 0.1 | 0.1 | 0.7 | 0.4 | 0.6 | 0.5 | 0.4 | >6.0* |
|  | Average recruitment rate (%): | 30.5 | 29.4 | 24.4 | 14.1 | 25.0 | 32.4 | 26.0 | <10.0 |
|  | Recruitment rate (min-max %): | 25.9–43.1 | 12.8–19.8 | 1.6–21.0 | 1.8–24.4 | 11.7–33.4 | 1.8–55.6 | 1.6–55.6 |  |
|  | Average annual 14-year population growth rate (%): | 7.8 | 12.3 | 11.6 | 5.9 | 10.4 | 9.7 | 9.6 | -10.2** |
|  | Last 5-year population growth rate (%): | 7.5 | 12.0 | 11.2 | 8.7 | 9.0 | 7.2 | 9.3 | -17.0** |
|  | Annual population growth rate (min-max %): | -28.8–21.5 | 6.8–19.8 | 2.7–21.1 | -5.0–10.9 | 5.1–19.1 | 0.6–19.6 | -28.8–21.5 |  |

\*Vital rates adapted from: [1] https://doi.org/10.3957/056.051.0100
\*\*Growth rate (Fig 9), period 2008 to 2021

### Performance of populations

Although cognisanze must be taken of the land size of the state-owned protected areas, such as that of the KNP (1.98 Mha) versus that of the non-state owned areas, and the subsequent difficulties reported to achieve operational efficiency in situational awareness, integrity management and access control, when comparisons of the rhino population sizes, population management and poaching rates on these smaller protected areas compared to that of the KNP, is done [55], the results show that provincial and private reserves do significantly better than the KNP itself under the current situation in South Africa (Table 4).

**Table 4.** SWR population sizes (Jan 2020) and poaching rates within four categories of land uses associated with rhino protection in South Africa during 2020.

|  | KNP | Other national parks | Provincial reserves | Private properties | Rhino properties excluding KNP | Governance index |
| --- | --- | --- | --- | --- | --- | --- |
| Nmber of white rhinos (n) = | 3,550<sup>1</sup> | 360 | 3,500<sup>2</sup> | 7,000<sup>3</sup> | 10,860 |  |
| Poaching rates: | 6.12% | 0.30% | 3.10% | 0.50% | 1.30% |  |
| Free State Province |  |  | 0.0% | 0.0% | 0.0% | 56.0% |
| Northern Cape Province |  | 0.0% | 0.0% | 0.015% | 0.009% | 58.6% |
| Eastern Cape Province |  | 0.0% | 0.0% | 0.0% | 0.0% | 67.7% |
| Gauteng Province |  |  | 0.0% | 0.029% | 0.018% | 82.2% |
| Western Cape Province |  | 0.0% | 0.0% | 0.015% | 0.009% | 87.6% |
| North West Province |  |  | 0.466% | 0.029% | 0.166% | 38.8% |
| Mpumalanga Province | 6.073% |  | 0.110% | 0.132% | 0.114% | 58.9% |
| Limpopo Province | 0.0047% | 0.30% | 0.027% | 0.250% | 0.175% | 65.0% |
| KwaZulu-Natal Province |  |  | 2.496% | 0.029% | 0.814% | 71.3% |

<sup>1</sup> numbers for KNP declined to 2,607 at end of 2020.
<sup>2</sup> numbers in Provincial reserves are most likely over-estimated.
<sup>3</sup> numbers on Private properties are most likely under-estimated.
Rates within provinces contribute relatively differently to countrywide poaching rates associated with the land uses.
Poaching incidences do not associate with an index of governance quality.
Data adapted from: [55], https://doi.org/10.17159/sajs.2021/11300

Data provided from private ownership for the six Case studies/ CBOs was analysed and the following vital rates assessed (Table 3) for comparison to data available for the “wild” SWR population in the KNP [1].

Natural annual population growth for white rhino is 8.0–12.0% per annum under ideal circumstances [6,33,56]. The average annual population growth performed by the management of the private rhino subpopulations since the first ownership in 1976 to 1987 (12 years) is 14.4% (Fig 2) with extreme deviations in the early years, and since 1988 to 2012 (25 years) is 8.6% (range 1.9–23.1%: end of 1^st^ quartile value 5.2%, end of 2^nd^ quartile value 6.7%, end of 3^rd^ quartile value 11.2%). The annual growth rate for the privately managed white rhinos accumulative (CBOs included) for 2013–2020 was 9.0%.

Annual biological growth performance achieved by the studied CBO Case studies respectively (Table 3 and Fig 8) measure as follow: **Case study no. 1** is 7.83% (average annual Recruitment Rate [RR] 30.46%, average annual Mortality Rate [MR] 1.43% and average annual Poaching Rate [PR] less than 0.10%) over 14 years; **Case study no. 2** is 12.34% (average annual RR 29.41%, average annual MR 0.86% and average annual PR less than 0.10%) over 9 years; **Case study no. 3** is 11.63% (average annual RR 24.41%, average annual MR 0.55% and average annual PR less than 0.70%) over 9 years; **Case study no. 4** is 5.87% (average annual RR 14.11%, average annual MR 0.25% and average annual PR less than 0.40%) over 11 years; **Case study no. 5** is 10.40% (average annual RR 25.0%, average annual MR 1.20% and average annual PR less than 0.60%) over 11 years; **Case study no. 6** is 9.73% (average annual RR 32.24%, average annual MR 1.20% and average annual PR less than 0.50%) over 12 years.

Average annual growth rate for the six white rhino CBO studies with *n* = 2,882 rhinos at the end of the study period, combined over 14 years is 9.6% (Table 3) despite being held semi-intensively, which is in line with the 9.0% (8.6% growth plus 0.4% off-take [4]), measured for the entire private white rhino population nationally (Fig 2). In addition, the CBO Case studies assessed, showed a loss of their populations due to poaching of less than 1.0% annually.

The annual growth rate for the KNP white rhino population calculated from the data in [1] was negative from 2008 to 2020 at minus −10.2% (range −26.4–2.3%: median −4.5%), in comparison with the overall positive trends (+9.6%) found for privately managed rhino populations (Fig 9).

**Fig 9.**
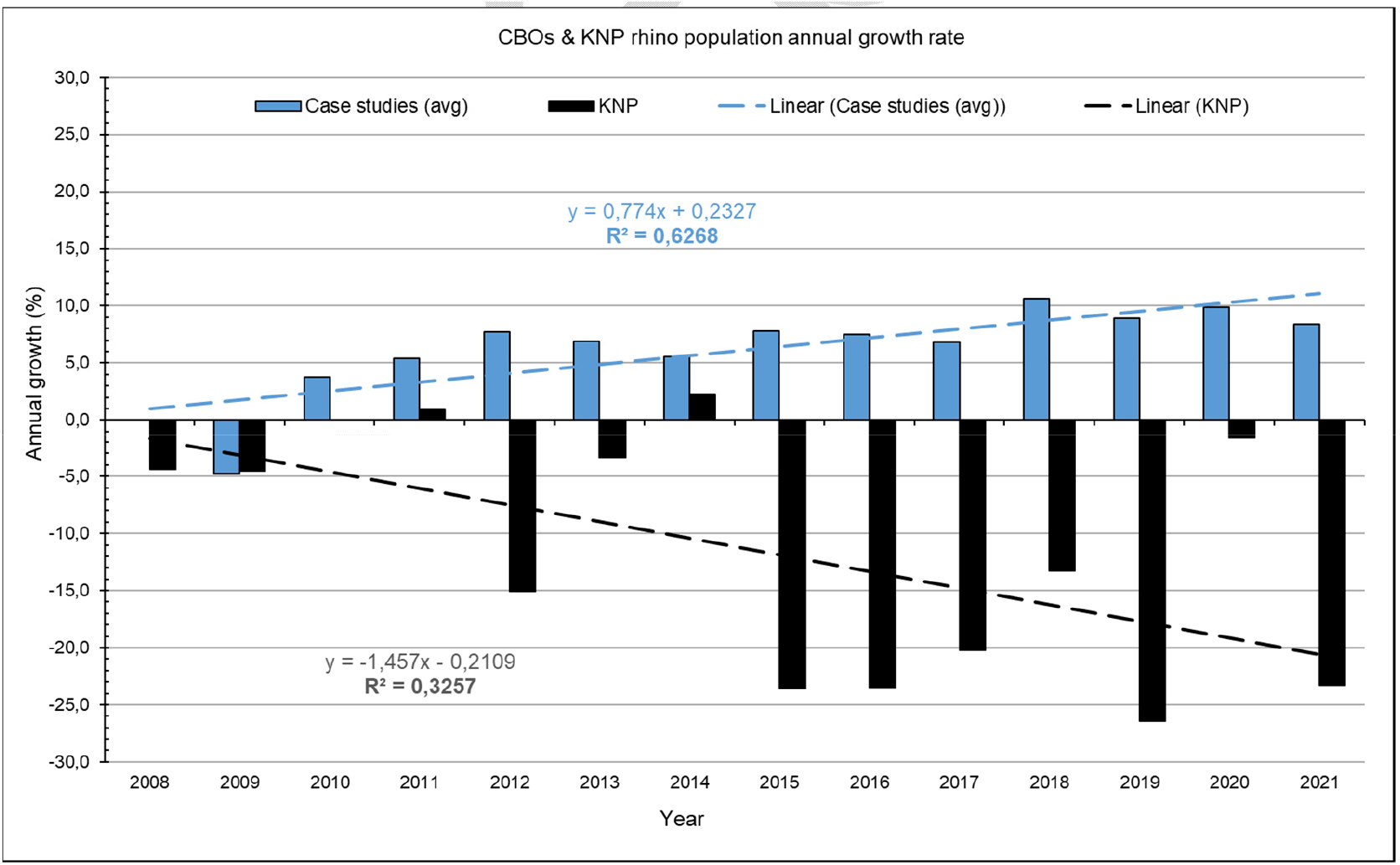
White rhino population trends since 2008, as for private CBOs and for the protected KNP. (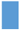 Upper curve) for CBO Case studies no’s 1–6 accumulative (*n* = 2,882 rhinos in 2021), +9.6% average annual increase in rhino numbers). (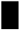 Lower curve) for KNP (*n* = 2,607 rhinos in 2021), −10.2% average annual decline in rhino numbers).

**Fig 10.**
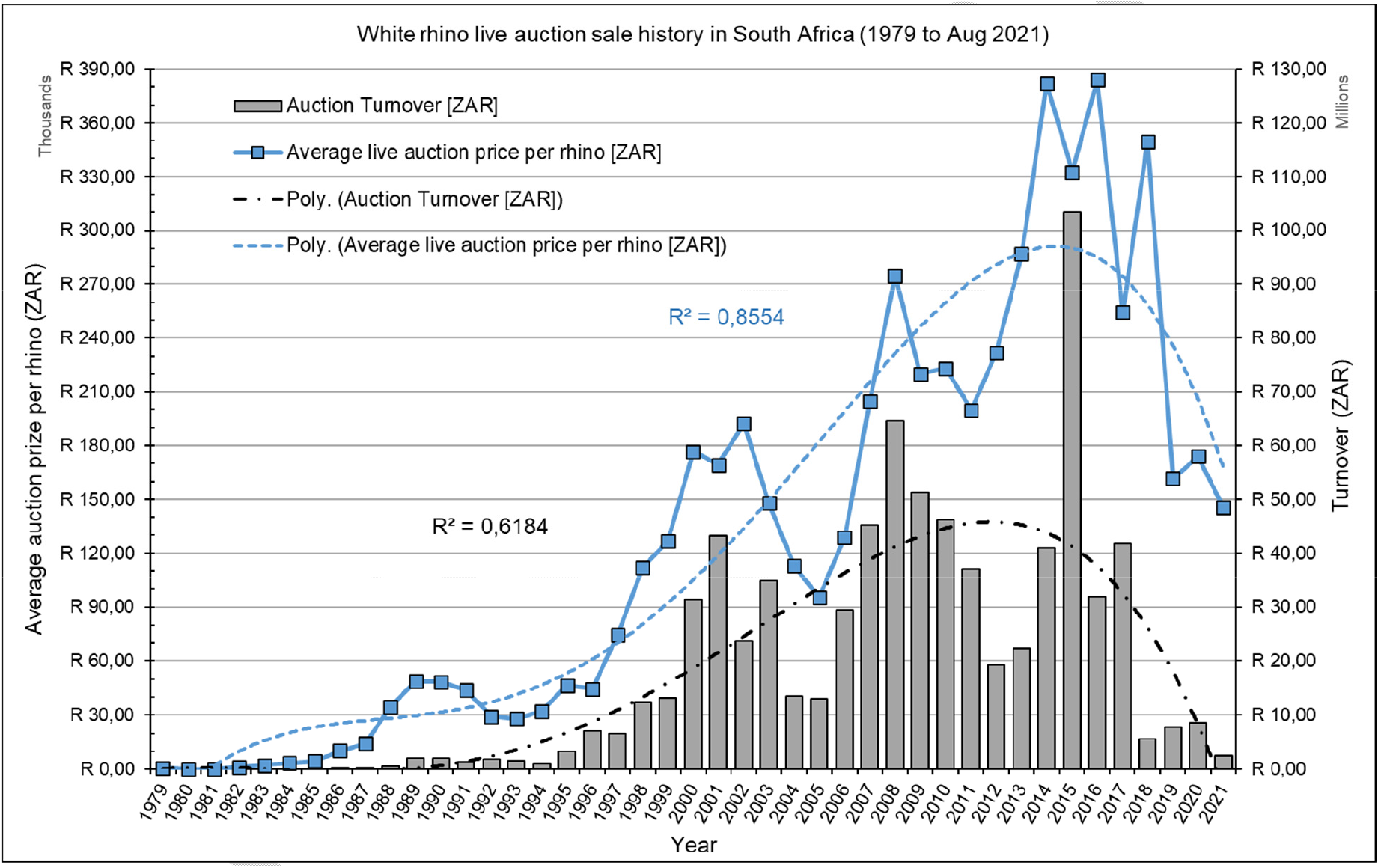
History of live auction sales value for white rhinos in South Africa. Data from Eloff and Cloete & Taljaard as cited in [49], Dr F Cloete, Head: Commercial Agri Finance, Agri Credit Solutions, South Africa, 28 Aug 2021.

## Discussion

This study emerged following the release of a High-Level Panel draft report (HLP, [27]) in May 2021 by the South African minister of DFFE proposing the possible closure and disbanding of South African private game ranching and disinvesting in captive breeding operations (CBOs) for southern white rhinoceros (SWR) populations. The *alleged* potential detrimental effect by the private sector to the conservation and sustainable survival of the species, as alluded to by the HLP draft report for rhinoceros, is in direct and significant contradiction to the findings of this study and representing the greater of the remaining global SWR species population.

### Species integrity, metapopulation management, national population

All extant southern white rhinos, irrespective of being managed as “wild”/ fenced-in state-protected populations, or as private and captive bred operations, are the progeny from the same single source, the same bottleneck gene-pool of less than <50 remaining animals from the former Mkuze Game Reserve (now HiMP) [4,6,24]. The current KNP white rhino population was re-introduced in October 1961 and again in 1987 from Mkuze. Since the SWR populations in the state-owned protected parks are predominantly enclosed and SANParks do not routinely introduce new rhino/ genes into the parks (an action refered to as metapopulation management), these populations are therefore potentially predisposed to the effect of genetic drift, loss of allelic diversity and inbreeding depression over time. All protected and wildlife entities in South Africa today (SANParks, provincial reserves, conservancies, wilderness heritage areas, trans-frontier parks, ranches, and farms) are fenced-in, fragmented, isolated systems (at different scale from <100 ha to 1.98 Mha) with restricted and/ or altered animal movement and different extend of reduced species genetics [48]. White, and black *Diceros bicornis* rhino, Lichtenstein’s hartebeest *Alcelaphus lichtensteinii*, oribi *Ourebia ourebi*, roan *Hippotragus equinus* and sable *H. niger* had been re-introduced to KNP as herds, some as small as *n* = 6 individuals per species from a single source from another fragmented and isolated (fenced-in) location. Most of these species had never been reinforced by new gene immigration/ gene-flow as to enhance its genetic integrity (heterozygosity).

To the contrary, the frequent commercial trade and subsequent translocations (gene-flow) between the private SWR subpopulation operations (game ranches), and especially since 1991 [6] with the legislation of the Game Theft Act and allowing for legal private ownership, fits the principle of metapopulation management and subsequent improvement of heterozygosity and species integrity [19,30,44–48]. Cumulative gene-flow for the six CBO Case studies accounts for metapopulation management between *n* = 176 different SWR subpopulations/ genetic pools, up to *n* = 24 and more different geographic sources per subpopulation. This gene flow narrative to be extrapolated with frequent trade and translocation between approximately *n* = 390 different private subpopulations in South Africa.

Notwithstanding the effects that the former genetic bottleneck holds for the species survival of the SWR, the health of global population numbers also needs be emphasized. The first survey of white rhino on private land as cited in [12] was undertaken in 1987 [57], and followed by many more surveys [**Error! Reference source not found.**,**Error! Reference source not found.**,39–40,43,58–62]. According to the latest published IUCN report [3], the African SWR population numbers were as follows for 2012 *n* = 21,320 (of which *n* = 18,933 occurred in South Africa); 2015 *n* = 20,560 (of which *n* = 17,208 [16,549–17,863: 90.0% CI] occurred in South Africa), 2017 *n* = 18,067 (17,212–18,915: 90.0% CI), of which *n* = 15,625 (90.0% CI) occurred in South Africa (Fig 1).

Examination of the listed poaching statistics of the supplementary information provided for this IUCN report, showed that of the *n* = 6,144 SWR reported as poached in South Africa during 2012–2017 (Fig 3), *n* = 5,864/6,144 (95.0%) were: [*n* = 643/668 (96.25%) in 2012, *n* = 966/1,004 (96.23%) in 2013, *n* = 1,161/1,215 (95.56%) in 2014, *n* = 1,113/1,175 (94.72%) in 2015, *n* = 1,011/1,054 (95.92%) in 2016, and *n* = 970/1,028 (94.36%) in 2017]. Unfortunately, no further subspecies breakdown of the annual poaching statistics were available at the time of this report but following the trend from the previous years, and taking an average of 95.0% of annual poaching losses being attributed to SWR for South Africa, the following estimates can be deduced, namely a total loss of: *n* = 769 (estimated *n* = 731 for a minimum 95.0% trend) in 2018, *n* = 594 (estimated at *n* = 564 for a minimum 95.0% trend) in 2019, and *n* = 394 (estimated *n* = 374 for a minimum 95.0% trend) in 2020. Therefore, totalling an estimated further loss of at least *n* = 1,669 SWR in South Africa alone due to poaching since 2017 to the end of 2020.

### Captive breeding operation (CBO)

CBO programs [63,114–115] are recognized as important components of conservation strategies for endangered/ threatened game species and has the potential to act as genetic and demographic reservoirs to reinforce wild populations [22]. In accordance with CITES Resolutions Conference 10.16 (Rev. CoP11, 2000): Captive bred refers to specimens or offspring born or otherwise produced in a controlled environment of parents that mated or otherwise transmitted their gametes in a controlled environment. In accordance with NEMBA and/ or TOPS, the following definitions apply: **(A) CBO** – means a facility where specimens of a listed threatened or protected animal species are bred in a controlled environment for conservation purposes; **(B) Bred in captivity** – means the offspring born or otherwise produced in a controlled environment of parents that mated or otherwise transmitted their gametes in a controlled environment, as described in CoP11; **(C) Controlled environment** – means a camp designed to hold specimens of a listed threatened or protected species in a way that, i) prevents them from escaping, ii) facilitates human intervention or manipulation in the form of the provision of food and water, artificial housing, or health care, and iii) facilitates the breeding or propagation of a listed threatened or protected species; **(D) Kept in captivity** – means that the species is kept in a controlled environment for a purpose other than transfer or transport, quarantine, or veterinary treatment; **(E) Bred in captivity** (as defined in Article I, paragraph (b) of the Convention, (CoP11) – means born or otherwise produced in a controlled environment and applied only, if parents mated or gametes were otherwise transferred in a controlled environment, or the parents were in a controlled environment when development of the offspring began; **(F) First-generation offspring (F1)** – are specimens produced from parents at least one of which was conceived in or taken from the “wild”; **(I) Second generation offspring (F2)** or subsequent generations (F3, F4 etc.) – are specimens produced in a controlled environment from parents that were also produced in a controlled environment.

As per the draft policy position document of DFFE [16, 27], the following important terms have not been defined, whereas Wildlife Ranching South Africa’s [64] definitions are as follows: **(G) Conservation** – quoted *“of elephants, lions, leopards, and rhinoceros is not about striking a balance between the well-being of people and wild animals. Conservation is about putting people first and managing all renewable natural resources, including big five wildlife, responsibility for the benefit of humanity. The responsible conservation management of elephant, lion, leopard, and rhinoceros must involve both ecological and economical sustainably for the socio-economic benefit of our people as supported by the South African Constitution”*. **(H) Conservation area management** – quoted *“is where wildlife is positively managed, produced and sustainably utilized for the socio-economic and environmental benefit of people. The creation of wealth from responsible or sustainable wildlife management, production and legal trade in wildlife and valuable wildlife products is promoted and facilitated through responsible conservation management. The conservation management of wildlife should not be confused with protected area or preservation management of wild animals as they have very different management objectives. Conservation management supports responsible sustainable use of all natural resources, does not prevent the use of wildlife by man but protects these resources against the abuse, pollution, erosion or their wasteful destruction”*. **(L) In protected or preservation management areas** – quoted *“wildlife is negatively protected, often at the expense of people, the sustainable use of these renewable natural resources is prohibited and the creation of wealth from the legal trade in wildlife and wildlife products is not facilitated and often denied”*. **(I) Economic sustainability** – quoted *“of a wildlife enterprise means that this business entity is economically self-sustainable, covers its own development and management costs and can survive, prosper or grow on its own ability to generate wealth.”*

According to the IUCN AfRSG and a later standard by the captive breeding community [18]: **(J) Semi-wild populations of rhinos** – occur mainly in small (<1,000 ha) areas, either in or out of the historical range of the taxon, and they live at a compressed density and spacing, requiring routine partial food supplementation and a high degree of management, but breed naturally; **(K) Captive populations of rhinos** – usually occur in small to very small areas (<100 ha), either in or out of the historical range of the taxon, and they have a compressed density and spacing, requiring partial or full food supplementation with frequent husbandry and veterinary intervention, and have a <u>manipulated breeding system</u>. In such situations rhinos may have controlled access to limited areas of natural habitat; **(L) Key 1 population** – population (*n*) increasing or stable and *n* >100 animals, also a population of rhinos whose survival is considered critical for the wider survival of the sub-species.

Important to note that the camp sizes of all subpopulations in the six CBO Case studies, as shown in Table 2, are greater than the maximum 100 ha cap defined for **captive populations (K)**.

### White rhinoceros species performance

The white rhinoceros is a relatively adaptable species able to survive in a variety of natural habitats from grassland to savanna and inhabit areas with mean annual rainfall ranging from 300 to 1,500 mm per annum. Juvenile mortality rates during the winter months on the Highveld are however high, which is thought to be due to a combination of low temperatures and poor grazing quality [50,65–66]. They do not appear to compensate for seasonal declines in food quality by switching to other fodder species or increasing the number of species eaten, and may instead draw on fat reserves during the dry season [67], or if possible, feed higher-up in the catena where reserve grazing of taller, but palatable decreaser grass species may occur. The poor response of white rhinos to temporarily adapt its feeding during drought or sparse grazing times often results in mortalities, especially in poorly managed protected areas, as experienced by the drought in 2015/ 2016 [26]. Better ecological management and provision of sufficient supplementary feeding on private white rhino CBOs contributes to sustaining an effective rhino population growth at an average of 9.0% per annum.

Inbreeding is generally prevented by the calves either leaving their mothers, or being pushed away by a breeding bull, when between the age of 2.5–3.5 years. The calves then join up with other adult females and/ or sub-adults, subsequently dispersing into new areas. Individuals of all ages have been recorded to disperse in search of better-quality graze and roam distances of up to 40–50 km during drought conditions [68–69]. Environmental barriers however may inhibit their dispersal. Males have non-overlapping territories which are known to range from 700–1,400 ha in typical savannas [70], and the boundaries of their home ranges are commonly aligned with topographic features such as rivers, watersheds, or roads [50]. Geographic translocation dispersal of white rhinos into multiple subpopulations by the private game industry play a vital role in overcoming the movement limitations set by natural and geographic barriers.

Analyses undertaken by the IUCN AfRSG indicated an average growth rate of the national white rhinoceros population of 7.1% from 1991 to 2012 [17,34,71], also see Fig 1. Several key events apparently contributed to a rapid increase since the all-time low *c*.1895, which include the development of chemical capture drugs, translocations, policy changes both locally and internationally, limited sustainable trophy hunting since 1961 [4,24], private ownership and live sale auctions. These have until recently created economic incentives for private ownership, thereby encouraging the expansion of rhinoceros range and numbers and significant numbers of white rhinoceros being translocated from “wild” populations to smaller secure areas where the animals are subjected to rewilded agro-sustainable biodiversity wildlife management and intensely protected against poaching. Poaching of privately owned white rhino being 10.0–20.0% compared to 60.0–80.0% in the government protected KNP. As stated by the NDF from the Scientific Authority [31], the future trend in the national population is unpredictable, and could increase by 1.9% or decrease by 3.9% after 5 years depending on the poaching scenario modelled. The −10.2% decline trend (Fig 9), and with a 75.0% loss since 2011 of the KNP population [1], emphasize global reality and the high importance of the privately managed rhino populations which grow positively at average 9.0% per annum (Fig 9), and despite a sustainable national annual hunting off-take quota of 0.4% [4].

### White rhino management and conservation narrative

The continuing loss of rhinoceros to poaching for their horn is currently the major threat to the survival of the species. Poaching increased each year from 2006, reaching a peak in 2014 when an estimated 6.5% of South Africa’s wild population was poached (Fig 3). In order for the current anti-poaching efforts to continue, significant financial inputs from external sources are required. The response to the threat has however been disproportionately high, removing much needed conservation funding from other species [27, 115].

In 2018, proportionally 72.0% [31] of the national white rhinoceros population was generally managed within state-protected fenced-in areas (52.0% in the fenced-in KNP), the national population with off-takes translocated in terms of ecological metapopulation management plans, mostly to private game ranches. A national white rhinoceros strategy was approved in 2000, and in December 2015 a national biodiversity management plan (BMP) for white rhinoceros was gazetted for implementation in terms of NEMBA. This plan was to form the basis for greater coordination between existing and future plans and is informed by the National Strategy for the safety and security of rhino populations in South Africa as well as the Rhinoceros Issues Management Report. The current HLP draft report [27] by its proposal to reserve investment of private rhino operations and 57.0% of the national white rhino population, deviates markedly from both the BMP as well as the findings of the Scientific Authority’s non-detriment finding (NDF) [31]. Since 2010, the South African government has launched a variety of initiatives in collaboration with various stakeholders to address the poaching threat and ensure the long-term conservation of the species, and in 2014 cabinet adopted an integrated four-pronged approach to curb rhinoceros poaching.

The data from the national rhino surveys referred to, and as shown in this study indicate most likely failure by government to secure the survival of its white rhinoceros population, whereas the data from the private rhino subpopulation operations indicate significant potential to save the southern white rhino from possible extinction, should government not proceed with the closure of as proposed by its HLP draft report.

SWR is listed as protected in terms of Section 56 of NEMBA and various provincial ordinances and acts provide further legislative protection. Permits are therefore required to undertake a variety of activities in relation to rhinoceros, e.g., hunting, keeping, selling and other forms of direct use. Legal off-take include management removals of animals and trophy hunting. An estimated 1.4% of the national herd is translocated live from state-protected areas annually [31], and mostly to private game ranching operations and CBOs. Between 2005 and 2016 a total of *n* = 774 live white rhinoceros were exported internationally from South Africa, 33.0% of the exports was for re-introduction purposes, 27.0% to zoos, and 26.0% to breeding facilities. The main destination countries were Namibia (38.0% of the exports), China (32.0%), and Botswana (7.0%).

#### Legal rhino trade

A moratorium to prohibit any sale of rhino horn or horn products within South Africa was implemented in February 2009. The moratorium was a temporary measure to afford DFFE an opportunity to develop and implement permanent measures aimed at eliminating the illegal international trade in rhino horn. The moratorium was set aside by the high court of South Africa in November 2015, thereby rendering the domestic trade in rhino horn within the borders of the country legal again. The amended norms and standards for the marking of rhinoceros and rhino horn and for trophy hunting purposes (published in April 2012) require that all rhino hunts are attended by conservation officials. In addition, the norms and standards require that an official must attend all dehorning activities. The regulations further require that a DNA sample must be collected from each animal, as well as from both horns, and all DNA samples are stored on the RhODIS database to ensure traceability. The system is well managed and rhino horn stockpiles are regularly audited.

As determined by [73] the mass of rhinoceros horn that could be obtained from CBO production varies from 5,319–13,356 kg per annum, and the amount of horn lost to poachers for the period 2012–2016 at 3,781–5,933 kg/year (average 5,718 kg per annum).

Currently live white rhino is listed as CITES Appendix II therefore may be traded, yet the inert horn is listed under Appendix I and may not be commercially traded. It makes no economic or conservation sense for an inert or non-living horn to have a higher CITES listing or protection than the living animal.

#### Poaching

Poaching, specifically in the KNP has disrupted the ability of SANParks to achieve its contribution to South Africa’s conservation targets for 2020 [1,3,31,35,73]. The counter reality of sustainable production wildlife management needed for the wealth of Africa’s people and poverty to fit the United Nations 2030 Agenda for Sustainable Development [80], are clearly emphasized and quantified by [74–**Error! Reference source not found.**] quote: *“Our planet is not to be conserved simply by locking away 30% by 2030 or 50% by 2050 from human use, but rather by managing 100% for sustainable human benefit”*.

Achieving targets such as annual growth rates of 5.0% is now deemed less feasible for white rhino “wild” populations in the current continual poaching onslaught [1,3,24,32,**Error! Reference source not found.**,76]. International growth rates for captive white rhino populations are similarly reported as showing negative annual growth rates (−3.5% as a percentage of the entire captive population).

Contrary to both the protected area populations as well as international intensive CBOs [33], the semi-intensive/ semi-wild private CBOs across South Africa has achieved average growth rates of 9.0% and has lost less than 2.0% of its populations to poaching per annum.

Trade bans, especially on ivory and rhino horn, has multiforcively indicated to fuel the fires of poaching, criminal syndicates, illegal or black-market trade, along with corruption associated with certain NGOs, politicians, and government officials. Trade bans are believed to be the cause of the poaching pandemic or the Rhino War [77].

#### Hunting

Legal and regulated hunting as assessed and argued by [4] has added positively to the sustainable-use conservation of this rhino species. The introduction of trophy hunting of adult white rhino bulls started in 1968 when there were only *n* = 1,800 animals.

It is estimated that the direct contribution of trophy and biltong hunters (all game species included) in 2016–2017 to the South African economy was ZAR 13.6 billion ($909 million), also the indirect and induced impact need be added [Van der Merwe 2018 as cited in 64]). Between 70.0% and 80.0% of trophy hunters’ spending takes place in the area of the hunt – in addition, hunting creates jobs, particularly in rural areas where employment is most needed. In three South African provinces by 2017 hunting created 31,500 jobs.

It was only after the first wildlife auctions in the late 1980s () that rhinos began to realise their commercial market value, incentivising private owners to also bring the rhino population up through breeding [78–79]. Rhino numbers and range then expanded considerably from this point. According to [24] the simultaneous development of more incentivising legislation (not by design!) around the ownership of wildlife saw the white rhino population grow to *n* >4,000 animals on approximately *n* = 400 private properties in South Africa by 2008 (Fig 2).

Prior to 2005 an average of 0.76% of the national white rhino population were hunted annually, from 2005 until 2012 it was 0.67% per annum, and since 2013 to present 0.40% per annum [4].

Trophy hunting selectively removes surplus and non-breeding adult males, whilst generating important revenue for private and state conservation (while poaching targets animals of all ages and sexes). Legal hunting, combined with the impact of poaching, has not yet reached a level where it has caused a cessation in population growth [4,24].

#### Private rhino ranching

Due to the significant economic benefits of hunting to game farmers (approximately $19 million over the period 2004–2008), together with live sales, the private sector has increasingly stocked white rhinos, effectively maintaining rapid meta-population growth, and contributing to the expansion of the species’ range by approximate 18,000 km^2^. The private sector in South Africa now conserves more rhinoceros than there are black and white rhinoceros in the whole of the rest of Africa. Live sales of surplus animals to the private sector have also been highly beneficial to conservation agencies, generating vital conservation revenue and preventing overstocking in established subpopulations.

In 2012 suggestions that South Africa would consider submitting a trade in rhinoceros horn proposal to the 17^th^ CoP to CITES saw an immediate temporary recovery in the average price for a white rhinoceros, [31].

Due to the increased rate of poaching, the cost of rhinoceros security has increased substantially in recent years. At the same time demand for trophy hunting has been declining while the international sale of legal rhinoceros horn remains prohibited under CITES. These factors have contributed to a negative shift in the cost benefit ratio of owning wild white rhinoceros and leading to a reduction in the live sale price since 2016 (). Income of the three largest rhinoceros sellers earned from the sale of white rhinoceros has reduced from a total of ~ZAR 100 million in 2009 when *n* = 370 rhinoceros were sold, to ZAR 20 million in 2014 when only *n* = 60 was sold. Furthermore, between 2009 and 2012 there was a reduction in the average price of white rhinoceros, from ZAR 365,000 per animal in 2009 to ZAR 258,000 in 2012, and less than ZAR 180,000 since 2019. Total loss of revenue from 2009 to 2012 alone is estimated at ZAR 373 million [31].

Reduced introduction of rhinoceros to new areas is expected to result in a significant decline in the metapopulation growth rate and the total population size, should the private rhino numbers be removed from the equation/ graph in Fig 1. A ripple-effect expected of loss of financial income to the conservation authorities that rely upon funds generated from rhino sales to the private sector to conserve rhinoceros in the state-protected areas, as outlined by the Scientific Authority in the rhino NDF [31].

The most important contribution made by privately held rhinoceros CBOs and related management practices, as described in the Case studies herein above to the positive conservation of rhinoceros in South Africa, and entirely for the owner’s own cost and personal risk, are still to be acknowledged both nationally and internationally.

It deserves to be mentioned again that the white rhino population at CBO Case study no. 1 is recognized as a KEY 1 population by the IUCN SSC’s AfRSG, and making a significant contribution to continental white rhino conservation. The CBO has successfully bred (F1), and (F2) generations, a total of *n* = 1,627 (as at end of May 2021) white rhino calves in a 14-year timespan.

#### Macro-genetic metapopulation species conservation management

Metapopulation management is a concept still needed to be defined for the uniqueness of the southern African context. Both practise and scientific studies indicate the private conservation model of frequent trading and translocation of individual animals across habitat and environmental boundaries, and between fragmented (fenced-in) subpopulations (which include SANParks, provincial and municipal protected areas, and private ranches), to have greatest positive effect restoring formerly bottle-necked and lost genetic integrity of most game species [48].

Recent studies of more than *n* = 4,000 buffalo *Syncerus caffer*, from *n* = 37 subpopulations [81–90] revealed the subpopulations from *n* = 22 private ranches to have greater genetic heterozygosity than the fenced-in KNP population. Thus, indicating enhancement obtained from metapopulation gene flow as a result from private live-trade and frequent translocation between private subpopulations. The same had been conquered through scientific research also for roan and sable [90], and ecologically for bontebok *Damaliscus pygargus pygargus* [91].

Further more, South Africa is the only country on the Continent after 2012 to 2018, to have managed to improve the genetic integrity of the national population of sable, as per number of hunted quality trophies entered in the Safari Club International (SCI) Trophy Record Register: *n* = 153 trophies from South Africa, *n* = 18 from Zambia, and less from the other countries. The South African trophies were mostly hunt on private game ranches [92–93], indicating the enhanced impact of private game ranching and fragmented metapopulation management restoring lost genetic integrity of exploited “wild” populations.

Multiple white rhino DNA tests recorded with Onderstepoort Veterinary Genetics Laboratory are still to be assessed, the outcome expected to conquer with that for the other game species above.

#### Biodiversity adaptation (environmental change)

Due to climate change the major of transformed land can never be restored to the previous pre-transformed biodiversity status, but can be rewilded to a different new, and improved agro-biodiversity wildlife status. Old, cultivated agriculture lands on private game ranches are rewilded to productive soil, which sustain a new but different floristic biodiversity. The new habitat supports a new and improved animal biodiversity, and contribute increased green economic production compared to the former cultivation [48,94–98]. The positive contribution of the private game sector to sustainable biodiversity and rhinoceros conservation are mostly misinterpreted and not recognised.

#### Global concept and management

An increase in rhino poaching since 2008 occurred despite the international ban on the commercial trade in rhino horn, and numerous law enforcement measures implemented in South Africa, which means that current protection measures of the protected area rhino populations have limited effectiveness. There is a certain economic value that could be derived from international trading of horn that are lost and could be allocated to rhinoceros protection [31].

As a result of the continued increase in the illegal trade in horn and the apparent failure of the CITES trade ban, there have been calls from various segments of the conservation community, as well as a plethora of peer-reviewed papers recently published in the scientific literature argue for a legal trade in rhinoceros horn [4,6,25,31,64,76,100–102]. Recommend that the CITES CoP seriously explore the merits of alternative regimes for rhinoceros horn trade, which involve more scope for legal trade than allowed under the presently applicable regime.

The white rhino NDF of the Scientific Authority [31] concluded: (a) that legal international trade in live rhinoceros to appropriate and acceptable destinations and the export of hunting trophies poses a low risk to the survival of the species; (b) that continued legal hunting is essential for the conservation and protection of rhino; (c) it is highly unlikely that the investment in the protection of rhinoceros emanating from current financial sources (government, external donors and private rhinoceros owners) can be sustained in the long term; (d) a legal trade in rhinoceros horn as an alternative source of funds should be explored; (e) the export of wild-sourced rhino horn for non-commercial purposes and horn sourced from captive breeding facilities that are registered in accordance with CITES Resolution Conf. 12.10 (Rev. CoP15) and meet the CBO criteria, as already approved by the Scientific Authority, will not be detrimental to the survival of the species in the wild. This conclusion contradicts the latest proposal by the minister [103] and the HLP draft report [27] of closure of the private rhino CBOs currently contributing to the protection of more than 60.0% of the global white rhino population.

The HLP report’s vision falls short of an achievable strategic objective for the conservation of the SWR. It focuses on the protection of these animals on only one land use option, rather than the positive management of agro-wildlife rewilding [2,48,104,115] for the socio-economic and environment benefit of our people on a variety of land use options that may be used for wildlife conservation [63].

South Africa is fast approaching the limit of available habitat for white rhino on state-owned land. This means that in order to continue to grow the species, new habitat within South Africa or the expansion of existing ranges in other states will be required soon.

The continued growth and expansion of the rhino populations and range through the introduction of herds in available new areas are therefore reliant on the private sector and communities making their land available for the introduction of rhinos sourced from protected areas and privately owned lands. The incentives for private landowners and communities to make their land available are mostly economic driven, including potential live sales from productive hers, eco-tourism and sustainable hunting. Due to the high levels of poaching and therefore the risks associated with the ownership of rhino the economic incentives are becoming outweigh by the costs relating to interventions required to secure rhino populations and has resulted in disinvestment of many rhino custodians as well as the establishment of more intensive protection zones and/ or captive breeding operations in South Africa, whereby a large number of rhino can be kept in areas small enough to provide more effective protection and security.

## Conclusions

The rhino population management dispute under the pressure of human growth, came in a new era of shift of thought and practise from **biodiversity conservation** towards **sustained-use conservation** with emphasis on species integrity [64,105–107]. The major of private rhinoceros CBOs in South Africa, as for the six Case studies herein above are rewilded [48,104] **agro-sustainable biodiversity wildlife conservation** systems either being semi-extensive/ semi-free roaming/ semi-wild, or semi-intensive, sustainable management.

This study found similar findings which further supports the notion that, rather than discounting the private sector and their successful contributions towards rhino conservation and promoting them to disband their rhino populations, they can provide key insight in alternative viable conservation strategies and management systems in response to the current southern white rhino’s plight for the survival and continued conservation. A key possible option would be to improve rhino security in the KNP by establishing rhino zones of reduced land size with higher rhino densities, similar to the populations found in the private reserves and that the “wild” rhino in the KNP could similarly benefit from these being in these intensive protective zones. A recent statement released by the DFFE on 8 February 2022, now suggests that SANParks is investigating the feasibility of additional actions such as establishing additional founder populations outside of the KNP, which is in direct contrast to the recommendations given by the Department in the HLP draft report [27].

Legal hunting of white rhinoceros has been beneficial to the conservation of the species in South Africa through expansion of its range and the maintenance of a rapid population growth, especially on private rewilded rangeland. Due to the significant economic benefits of hunting to game ranchers [4], together with live sales and ecotourism, the private sector has increasingly stocked these animals. This has contributed to the expansion of the species’ range and has maintained a positive annual population growth of 9.0%, though still slightly short to the nett balancing of the minus - 10.20% annual decline suffered with the KNP population. However, the current prohibition on the commercial international trade in rhino horn can be viewed as a missed opportunity for beneficiation of owning and protecting rhinoceros [31]. Privately owned game farms have contributed significantly to white rhinoceros conservation, with approximately 57.0% of the national herd (more than >7,000 animals in 2020, currently projected at >8,000 [R^2^ = 0,988]) kept on approximately 18,000 km^2^ of privately owned land.

Southern white rhinoceros as a species is conservation dependent, occurring solely in protected areas and on private game ranches, but it is tolerant to local human activity and can be ranched under rewilded semi-intensive and semi-extensive conditions. Under these conditions, where the density of animals is higher and regular anaesthetic procedures for management purposes and/ or translocation are likely to increase stress levels, there is no detectable difference in cow fertility [108–109]. Also, faecal glucocorticoid metabolite (fGCM) levels do not differ between ranched and free-ranging adult rhino individuals, but routine dehorning procedures do result in short-term stress responses that dissipate after 72 hours [110].

Despite CBO registrations and macro-genetic metapopulation genetic diversification through gene migration/ gene flow by way of frequent translocations and relocations [19,22,44–48] across more than *n* = 390 privately managed white rhino populations (Fig 2), the HLP draft report [27] insubstantially (with no scientific support) claims these privately ranched animals to be genetically inferior and of little to no value to future conservation and survival of the species.

However, summarised parameters from the six CBO Case studies are as follow:

1. These CBOs house in total *n* = 2,882 (2020–2021 as per end date of respective study) white rhinos which is 20.6% of the estimated working national South African white rhino population as of the end of 2020, inhabiting a total land area size of 22,762 ha.
2. Average (large) camp size per breeding subpopulation is 749 ha (range 100–1,605 ha), the minimum average natural territory size for “wild” white rhinos is 700 ha.
3. In total 97.6% of the land inhabited by these CBOs are rewilded (20.4% from previously ploughed crop land, and 82.9% from previously alien and/ or mono culture livestock farming).
4. Five of the six rhino CBOs studied shared its natural habitat with other multi-species wildlife.
5. All six operations studied has the following best practise ecological strategies in common:

a. semi-free roaming systems, with partial supplement feeding for optimal health and reproduction;
b. *n* = 2–10 adult breeding bulls per subpopulation, allowing breeding females self-choice of natural selection of mating partners;
c. *n* = 3–5 or more sub-adult bulls in each subpopulation to allow for the natural establishment of social hierarchy;
d. ecological veld management either by: (i) rotational grazing of breeding herd across 2–4 sister camps, or (ii) a programme of multiple fire/ burn blocks per camp;
e. maintain average population growth rates around 9.6%, compared to <2.0% for most protected parks, and minus −10.2% for KNP (international IUCN goal for wild populations is 5.0%, and rarely achieved anymore);
f. a founder stock sourced from *n* = 9–95 different external genetic resources (populations) per CBO respectively, enhancing macro-genetic metapopulation gene flow, compared to KNP population that is limited to only one genetic source;
g. four of the six operations studied periodically had to extend its land area by purchasing additional land at between 1.5 and 3.5 million ZAR per 100 ha (an average of 400–800 ha needed for each new subpopulation) for making space for progeny growth – this is done without being allowed to generate income from legal trade of its stockpile of rhino horn produced;
h. all Case studies are burdened by excessive costs for security measures and management;
i. all Case studies are either semi-intensive or semi-extensive protected rewilded systems, and of major difference from any zoo, safari park, or canned system.

Performance as measured with private rhino populations in South Africa disclaim the argument of species diversity demise [111] of the SWR, rather than greater geographic metapopulation species diversity sustaining gene flow and improved species integrity. The study provides significant evidence for mind-shift of DFFE and IUCN towards agro-sustainable-use biodiversity conservation through private macro-genetic metapopulation management of rhino since 1961.

Community Based Conservation (CBC) another approach propagated by the HLP draft report [27] – Can it or does it work? The answer given by Professor Marshall Murphree of the University of Zimbabwe, and a principal architect of the world-famous CAMPFIRE program in Zimbabwe, as cited by [112], quoted answer: *“Sometimes it does, and sometimes it doesn’t.”* So the better question to ask is this*: “Under what does it deliver conservation goals as well as community benefits?”* There is no single answer to this question – quote: *“Conservation is no longer the application of biological science – it comprises many cross-cutting elements and interdisciplinary themes. The science of ecology is not sufficient to describe its limits. Now that people are considered to be part of ecosystems, and not simply the external managers, we must now speak of political ecology and environmental economics.”* Furthermore, the well-known Dr Ian Player, who more than anyone else was responsible for making the SWR safe by its wider distribution, also was a sincere advocate of its **sustainable utilisation as a strategy for its survival**, and a strong supporter of a legal trade in rhino horn [113].

It has been postulated that small or isolated populations, like those kept in zoos or safari Parks, are vulnerable to reproductive difficulties and may suffer from biased sex ratios, increased inbreeding, and fluctuations in reproductive successes. No reproductive difficulties had been noted from the accumulative 14-years of practice and management of the white rhinos in the six Case studies.

## Author Contributions

**Conceptualization:** Deon Furstenburg, Michelle Otto, Derek Lewitton.

**Data curation:** Deon Furstenburg, Michelle Otto, Pieter v Niekerk, Derek Lewitton.

**Formal analysis:** Deon Furstenburg, Michelle Otto.

**Funding acquisition:** Deon Furstenburg, Derek Lewitton.

**Resources:** Deon Furstenburg, Michelle Otto, Pieter v Niekerk.

**Supervision:** Deon Furstenburg.

**Writing ± original draft:** Deon Furstenburg, Michelle Otto.

**Writing ± review & editing:** Michelle Otto, Deon Furstenburg, David Balfour

## Notes

### Competing Interest Statement

The authors have declared no competing interest.

